# Single-cell atlas reveals alveolar macrophage-specific Oncostatin-M drives pulmonary fibrosis via fibroblast activation in PCV2d infection

**DOI:** 10.64898/2025.12.08.692949

**Authors:** Jingmei Xu, Xinyi Fan, Panlong Chen, Yumei Yuan, Yunlong Chen, Gang Fan, Bin Yang, Fan Yang, Yan Wang, Shiqiang Zhang

## Abstract

A hallmark of pulmonary fibrosis is the aberrant activation of lung fibroblasts into pathological fibroblasts that produce excessive extracellular matrix(1). Clarifying the cell-specific mechanisms by which viruses induce this process is crucial for developing effective strategies to intervene in disease progression. Here, we demonstrate that alveolar macrophage subsets specifically express tumor suppressor M (Osm) after infection by PCV2d infection and related intervention experiments in mice, combined with single-cell transcriptome sequencing (scRNA-seq). Through validation using histopathological analyses and an in vitro macrophage-fibroblast co-culture model, these alveolar macrophage-derived oncostatin M (Osm) were confirmed to specifically drive fibroblast activation. Vaccine intervention experiments further confirmed that targeted inhibition of oncostatin M (Osm) expression in alveolar macrophages significantly attenuated fibroblast activation and extracellular matrix deposition, thereby mitigating pulmonary fibrosis progression. Therefore, subgroup-specific secretion of oncostatin M (Osm) by alveolar macrophages represents a key driver mediating PCV2d infection-induced pulmonary fibrosis, and targeting this pathway offers a potential therapeutic strategy for viral-induced pulmonary fibrosis.

## Introduction

Pulmonary fibrosis (PF) is a life-threatening interstitial lung disease characterized by progressive extracellular matrix deposition and irreversible impairment of lung function(2). Currently, clinical therapeutic options for this condition remain extremely limited. Accumulating evidence has demonstrated that respiratory viral infections, such as influenza virus and SARS-CoV-2, can induce pulmonary fibrosis;however, the cell-type-specific mechanisms underlying virus-fibroblast crosstalk remain poorly elucidated.Although oncostatin M (Osm) secreted by alveolar macrophages has been reported to activate fibroblasts via the JAK-STAT pathway(3), whether Osm is specifically derived from a subset of alveolar macrophages and its precise regulatory role in viral-induced pulmonary fibrosis remain unknown.

The porcine circovirus type 2d (PCV2d) infection model offers a unique opportunity to dissect these mechanisms: this virus induces interstitial pneumonia and progressive pulmonary fibrosis in mice, and its strong tropism for pulmonary macrophages provides an ideal platform for investigating cellular heterogeneity.Crucially, current research is generally restricted to bulk cell analyses, lacking investigations into the spatiotemporal dynamics of Osm secretion by specific alveolar macrophage subsets, and has failed to establish a causal link between this pathway and fibrosis progression.

This study integrated single-cell RNA sequencing (scRNA-seq), histopathological analyses, and macrophage-fibroblast co-culture models to identify the specific cellular sources of oncostatin M (Osm) in PCV2d-infected lungs. Herein, we demonstrate for the first time that PCV2d infection selectively activates subgroup-specific overexpression of Osm in alveolar macrophages, which in turn drives fibroblast activation and collagen deposition, ultimately contributing to pulmonary fibrosis.This mechanism was validated using scRNA-seq trajectory analysis, in vitro co-culture assays, and PCV2d vaccine intervention experiments, thereby laying a theoretical foundation for anti-fibrotic strategies targeting Osm-producing alveolar macrophages.

## Materials and methods

### Pathogenicity studies in mice

To study the pathogenicity of the isolates in mammals, the 6th-week-old healthy BALB/c mice were selected for infection test. Mice were purchased from the Chengdu Dossy Experimental Animals CO. (Chengdu, China) and we promise that the study was performed according to the international, national and institutional rules considering animal experiments, clinical studies and biodiversity rights. The study protocol was approved by the Institutional Animal Care and Use Committee of Northwest A&F University(approval number: XN2023-0903).

### Virus strain

In 2023, this study collected lymphatic tissue samples from piglets suffering from respiratory diseases and emaciation symptoms from a large-scale pig farm in Shandong Province. One gram of sample was added to 1×PBS solution containing 200 IU/mL penicillin and 0.2 mg/mL streptomycin. After grinding, the sample underwent three freeze-thaw cycles, followed by centrifugation at 10,000 rpm for 30 minutes. The supernatant was then filter-sterilised through a 0.22 μm pore membrane. Five hundred microlitres of the filter-sterilised ground inoculum was inoculated into PK-15 cells with a 70-80% confluence (cultured in a T25 flask) and adsorbed for 2 hours in a 37°C, 5% CO2 incubator, with shaking every 30 minutes. After adsorption, the supernatant was discarded. Subsequently, 5 mL of DMEM medium (Dulbecco’s Modified Eagle Medium, Gibco, 2393822) containing 2% FBS (heat-inactivated foetal bovine serum, LONSERA, S711-001) was added, and cultivation continued. After 72 hours of cultivation, the cells underwent three freeze-thaw cycles, and the supernatant was collected by centrifugation to obtain the F1 generation of the virus. Five hundred microlitres of the F1 generation virus was inoculated into PK-15 cells and passaged for five consecutive generations, followed by five rounds of blind passage without conditions. Subsequently, the cells were inoculated into the PK-15 cell line and passaged for another five generations, and finally, the culture was preserved in a -80°C environment.The sequence of the PCV2d strain was submitted to NCBI, with the GenBank accession number OQ730503.

### Animal model construction

Mice were randomly divided into 4 groups,with 3 mice in each group: the PCV2d-infected group (1×105.14 TCID50 per mouse). Control mice were infected with PBS. The adjuvant control group (Adjuvant group, intramuscular injection with aluminium salt adjuvant). The vaccine-treated group (Vaccine group, intramuscular injection with inactivated PCV2d vaccine). On day 30 post-immunisation, all mice in the adjuvant control group and the vaccine-treated group were intraperitoneally injected with PCV2d virus (1×105.14 TCID50 per mouse). On day 14 post-infection, mice were euthanised by cervical dislocation, and lung tissues were collected for subsequent analysis.

### Sample preparation and observation

Five healthy Balb/c mice aged 6 weeks were subcutaneously inoculated with SD02 inactivated vaccine at 3 times dose (1 mL), and another 3 mice were taken as blank control. After 10 days of observation, liver, spleen, lung and kidney tissues of mice in each group were fixed in 4% polyformaldehyde solution and sent to Wuhan Saiwei Biological Company for HE staining sections for histological observation.

HE staining was used to observe the changes of lung tissue structure under light microscope. ImageJ software was used to quantitatively analyze the alveolar vacuolation area of 3 fields randomly selected from each section, and the percentage of vacuolation area in the total area of field was calculated.

### Tissue dissociation and preparation

The fresh tissues were stored in the sCelLiveTM Tissue Preservation Solution (Singleron) on ice after the surgery within 30 mins. The specimens were washed with Hanks Balanced Salt Solution (HBSS) for three times, minced into small pieces, and then digested with 3 mL sCelLiveTM Tissue Dissociation Solution (Singleron) by Singleron PythoN™ Tissue Dissociation System at 37 °C for 15 min. The cell suspension was collected and filtered through a 40-micron sterile strainer. Afterwards, the GEXSCOPE® red blood cell lysis buffer (RCLB, Singleron) was added, and the mixture[Cell: RCLB=1:2 (volume ratio)] was incubated at room temperature for 5-8 min to remove red blood cells. The mixture was then centrifuged at 300 × g 4 ℃ for 5 mins to remove supernatant and suspended softly with PBS.

### RT & Amplification & Library Construction

Single-cell suspensions (2×105 cells/mL) with PBS (HyClone) were loaded onto microwell chip using the Singleron Matrix® Single Cell Processing System. Barcoding Beads are subsequently collected from the microwell chip, followed by reverse transcription of the mRNA captured by the Barcoding Beads and to obtain cDNA, and PCR amplification. The amplified cDNA is then fragmented and ligated with sequencing adapters. The scRNA-seq libraries were constructed according to the protocol of the GEXSCOPE® Single Cell RNA Library Kits (Singleron)(4).Individual libraries were diluted to 4 nM, pooled, and sequenced on Illumina novaseq 6000 with 150 bp paired end reads.

### Primary analysis of raw read data

Raw reads from scRNA-seq were processed to generate gene expression matrixes using CeleScope (https://github.com/singleron-RD/CeleScope) v1.9.0 pipeline. Briefly, raw reads were first processed with CeleScope to remove low quality reads with Cutadapt v1.17 to trim poly-A tail and adapter sequences. Cell barcode and UMI were extracted. After that, we used STAR v2.6.1a(5) to map reads to the reference. UMI counts and gene counts of each cell were acquired with featureCounts v2.0.1(6) software, and used to generate expression matrix files for subsequent analysis.

### Quality control, dimension-reduction and clustering (Seurat)

We used functions from Seurat v3.1.2(7) for dimension-reduction and clustering. Then we used NormalizeData and ScaleData functions to normalize and scale all gene expression, and selected the top 2000 variable genes with FindVariableFeautres function for PCA analysis. Using the top 20 principle components, we separated cells into multiple clusters with FindClusters. Batch effect between samples was removed by Harmony(8). Finally, UMAP algorithm was applied to visualize cells in a two-dimensional space.

### Differentially expressed genes (DEGs) analysis

To identify differentially expressed genes (DEGs), we used the Seurat FindMarkers function based on Wilcox likelihood-ratio test with default parameters, and selected the genes expressed in more than 10% of the cells in a cluster and with an average log(Fold Change) value greater than 0.25 as DEGs. For the cell type annotation of each cluster, we combined the expression of canonical markers found in the DEGs with knowledge from literatures, and displayed the expression of markers of each cell type with heatmaps/dot plots/violin plots that were generated with Seurat DoHeatmap/DotPlot/Vlnplot function. Doublet cells were identified as expressing markers for different cell types, and removed manually.

### Pathway enrichment analysis

To investigate the potential functions of DEGs, the Gene Ontology (GO) and Kyoto Encyclopedia of Genes and Genomes (KEGG) analysis were used with the “clusterProfiler” R package 3.16.1(9). Pathways with p_adj value less than 0.05 were considered as significantly enriched.

### Cell type annotation

The cell type identity of each cluster was determined with the expression of canonical markers found in the DEGs using SynEcoSys database. Heatmaps/dot plots/violin plots displaying the expression of markers used to identify each cell type were generated by Seurat v3.1.2 DoHeatmap/DotPlot/Vlnplot.

### In vitro co-culture system

Porcine alveolar macrophages (PAM) were isolated from lung lavage fluid of healthy piglets, purified by adherent method and cultured in RPMI 1640 medium at 37℃ and 5% CO2. Porcine lung fibroblasts (PLF) were obtained from piglet lung tissue by tissue mass method and cultured in DMEM medium at 37℃ and 5% CO2.

In order to verify the feasibility of the key regulatory mechanism of pulmonary fibrosis in the cross-species model, we constructed a heterologous co-culture system of porcine macrophages and lung fibroblasts. PAM and PLF were seeded in 24-well plates at a ratio of 1:1. After 24 hours of culture, the cells were attached and infected with PCV2d (MOI=1). After 24h of infection, cells were harvested, total RNA was extracted, and used for qRT-PCR assay.

### Real-time quantitative PCR (qRT-PCR)

Total RNA was extracted from cells or tissues using TRIzol reagent (Invitrogen) and RNA concentration and purity were determined by NanoDrop 2000 to ensure OD260/280 was between 1.8 and 2.0. The extracted mRNA was reverse-transcribed using the reverse transcription reagent in PerfectStart® Green qPCR SuperMix. RT-qPCR reactions were performed using PerfectStart® Green qPCR SuperMix reagent and qPCR reactions were performed using BioRad real-time quantitative PCR instrument.

All quantitative data in this assay were obtained using Bio Rad CFX96 real-time fluorescence quantitative analyzer to obtain original Ct values. The relative expression of qPCR was calculated using qPCR relative quantification calculation method--2^-(<$Ct). GraphPrism8.3.0 software was used for statistical analysis and mapping of RT-qPCR and other test data. The test data came from three independent replicates, and the statistical results were presented in the form of “mean ± standard deviation”. For one-way test, the comparison of means between two groups of independent samples was performed by Student’s t-test, and the comparison of means between two groups of samples was performed by Dunnett’s multiple comparisons test. Specifically: *P<0.05, **P<0.01, ***P<0.001, ***P<0.0001, ns means no significant difference.

## Results

### 1. Histopathological Phenotype Analysis of PCV2d Infected Lung Fibrosis

To explore the role of PCV2d infection in pulmonary fibrosis, we established a murine model. Mice were intraperitoneally injected with either PBS (MOCK group) or PCV2d virus (Fig 1A). Lung tissues were harvested 30 days post-infection for histopathological analysis and single-cell RNA sequencing (scRNA-seq), aiming to identify the histological changes and alterations in pulmonary cell subsets induced by the infection.

**Figure 1.**
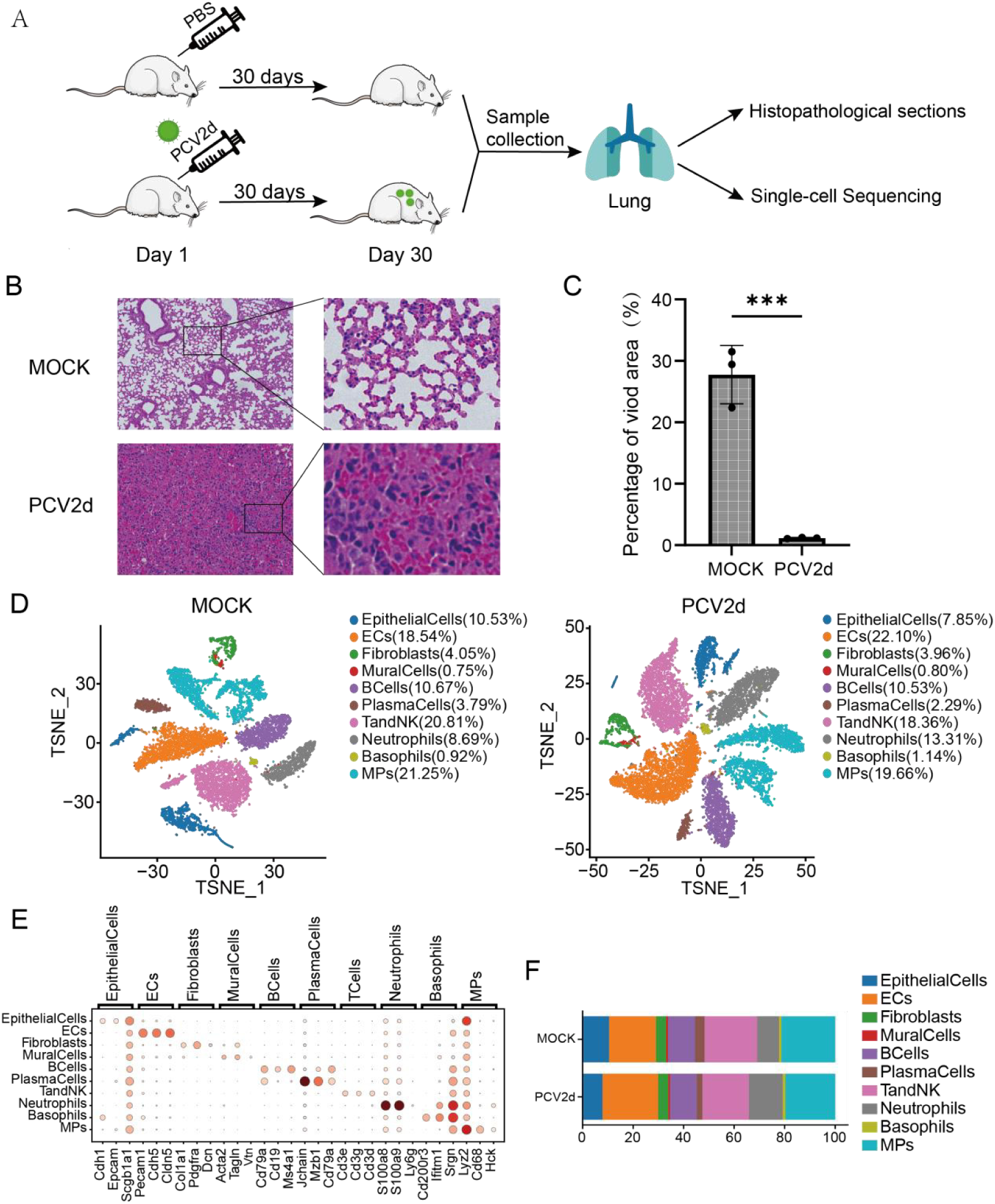
Design and differences in lung cell composition investigated throughout PCV2d infection in BALB/c mice. (A) Test flow design. (B) Differences in lung fibrosis sections between MOCK group and PCV2d group. (C) Count the percentage of vacuoles in three random visual fields. (D) Cell grouping in both treatment groups. (E) Subpopulation difference marker gene. (F) Percentage of each subgroup in both treatment groups.

Thirty days after PCV2d infection, the lung tissues of mice exhibited notable alveolar architectural disruption and interstitial widening (Fig 1B). ImageJ analysis showed that the percentage of alveolar vacuolization area in the infected group was significantly lower than that in the control group (Fig 1C). Using scRNA-seq technology, 54,771 cells isolated from murine lung tissues were sequenced for 11 cell types (Fig 1D). The experimental data were visualized via the T-SNE method (Fig 1E), encompassing multiple cell types such as epithelial cells, endothelial cells, fibroblasts, mural cells, B lymphocytes, plasmablasts, T and NK cells, neutrophils, basophils, and mononuclear phagocytes.Among these, the proportion of mononuclear phagocytes (MPs) was significantly decreased, while the proportion of neutrophils was markedly increased. Additionally, the proportion of endothelial cells (ECs) also rose (Fig 1F). Given the relatively low abundance of proliferating cells, they were not analyzed in this study.

PCV2d infection induced typical pathological phenotypes of pulmonary fibrosis, manifested as alveolar architectural destruction, interstitial widening, and a substantial reduction in the alveolar vacuolization area. Single-cell analysis further revealed infection-specific changes in the pulmonary immune microenvironment: a downregulation of MPs and an expansion of neutrophils, indicating that the reprogramming of myeloid cell subsets is involved in the fibrotic process.

### 2. Expression characteristics of pulmonary fibrosis-related genes Col1a2 and Timp3 in lung fibroblasts from infected and healthy groups

To evaluate the impact of PCV2d infection on the activation of pulmonary fibroblasts, this study employed single-cell RNA sequencing (scRNA-seq) to systematically compare the transcriptional characteristics of 1008 fibroblasts in the lung tissues of the MOCK group and the infection group (Fig 2A), among which 449 cells were from the MOCK group and 559 cells were from the infection group. The cells were visualized using the TSNE method, and the results are shown in (Fig 2B). Compared with the MOCK group, the expression levels of Col1a2 (encoding type I collagen) and Timp3 (matrix metalloproteinase inhibitor) were significantly increased in the infection group (Fig 2C). Further validation through T-SNE visualization analysis revealed a significant increase in the proportion of Col1a2 and Timp3 positive cells in the infection group (Fig 2B). We classified the fibroblasts into four subgroups through clustering methods (Fig 2D). Among them, the Fibroblasts_1 subgroup simultaneously highly expressed Col1a2 and Timp3 (Figs 2E and 2F), and the proportion of this subgroup significantly increased in the infection group. Differential gene analysis showed that the Fibroblasts_1 subgroup specifically expressed myofibroblast activation markers (such as Acta2) (Fig 2G).

**Figure 2.**
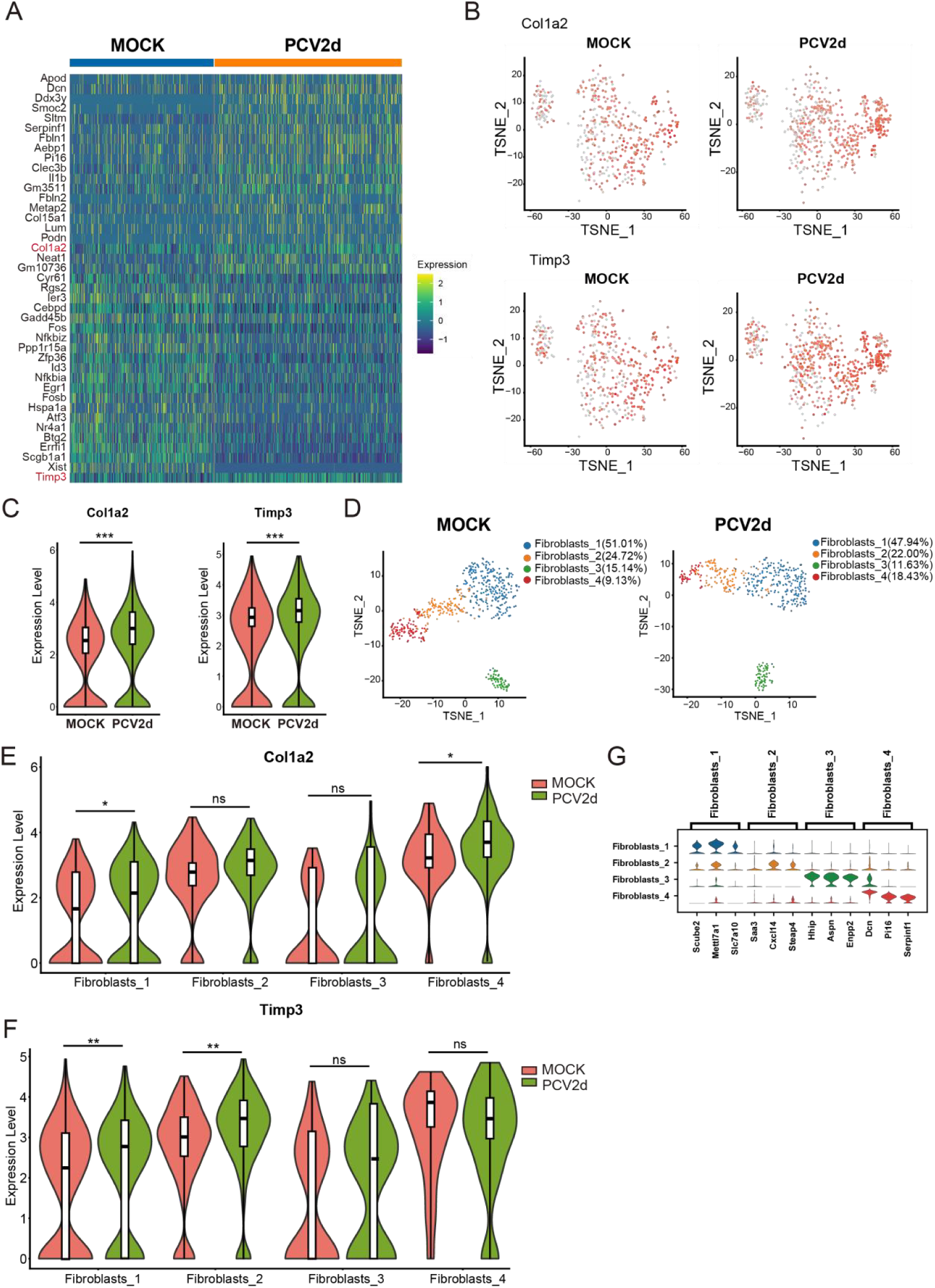
Analysis of fibrosis factors in MOCK group and PCV2d infection group in lung fibrosis phenotype caused by virus infection. (A) Total differential genes in fibroblasts under two treatments. (B) Fibroblast factors COL1A2 and TIMP3 were expressed in the fibroblast population of both treatment groups. (C) Violin expression levels of COL1A2 and TIMP3 genes infibroblasts under bothtreatment groups. (D) Secondary grouping of fibroblasts under both treatment groups. (E) Violin expression of COL1A2 in each subgroup under both treatment groups. (F) Violin expression of TIMP3 in each subgroup under two treatment groups (G) Marker gene in four subgroups.

In conclusion, PCV2d infection can specifically activate the pro-fibrotic phenotype of pulmonary fibroblasts, with the main feature being the significant upregulation of Col1a2 and Timp3 expression. Through subgroup analysis, it was further clarified that the Fibroblasts_1 subgroup is the key effector population involved in this process. This subgroup simultaneously highly expresses genes related to collagen synthesis and matrix remodeling, suggesting its core role in mediating the abnormal deposition of extracellular matrix (ECM).

### 3. Screening of Key Inflammatory Factors in Viral Infection Induced Pulmonary Fibrosis

Pulmonary fibrosis occurs as a consequence of many types of severe lung injury and is largely associated with inflammatory responses(10). Pulmonary fibroblasts are the core effector cells in the pathological process of pulmonary fibrosis(11), and their functional activation is finely regulated by various cell types in the microenvironment. In this study, we identified nine cell subsets with key regulatory functions in the microenvironment of fibrotic lung tissue. These cells release specific inflammatory factors through paracrine mechanisms to induce the transformation of pulmonary fibroblasts to a pro-fibrotic phenotype.

To systematically dissect this intricate intercellular crosstalk network, the present study employed an integrated multi-omics analysis strategy. Initially, differential gene expression profiles across nine distinct cell types were comprehensively analyzed to identify common genes with significant expression changes across multiple cell populations (Figs 3A-I).This step aimed to pinpoint conserved molecular signatures that might underpin the coordinated cellular responses in the context of fibrosis.Subsequently, these candidate genes were subjected to cross-validation against a curated database encompassing 17 well-documented pro-fibrotic inflammatory factors, as reported in the existing literature(12). This critical filtering step served to prioritize genes with established links to fibrotic processes, thereby enhancing the biological relevance of the findings. Through rigorous bioinformatic validation and functional enrichment analysis, interleukin-1B (IL1b) and oncostatin M (Osm) emerged as the core regulatory factors that not only displayed consistent differential expression across the majority of the analyzed cell types but also demonstrated robust associations with the known pro-fibrotic inflammatory pathways in the validation database. Their convergence in both the transcriptomic profiling and the literature-based validation solidified their roles as key mediators in orchestrating the fibrotic cascade.

**Figure 3.**
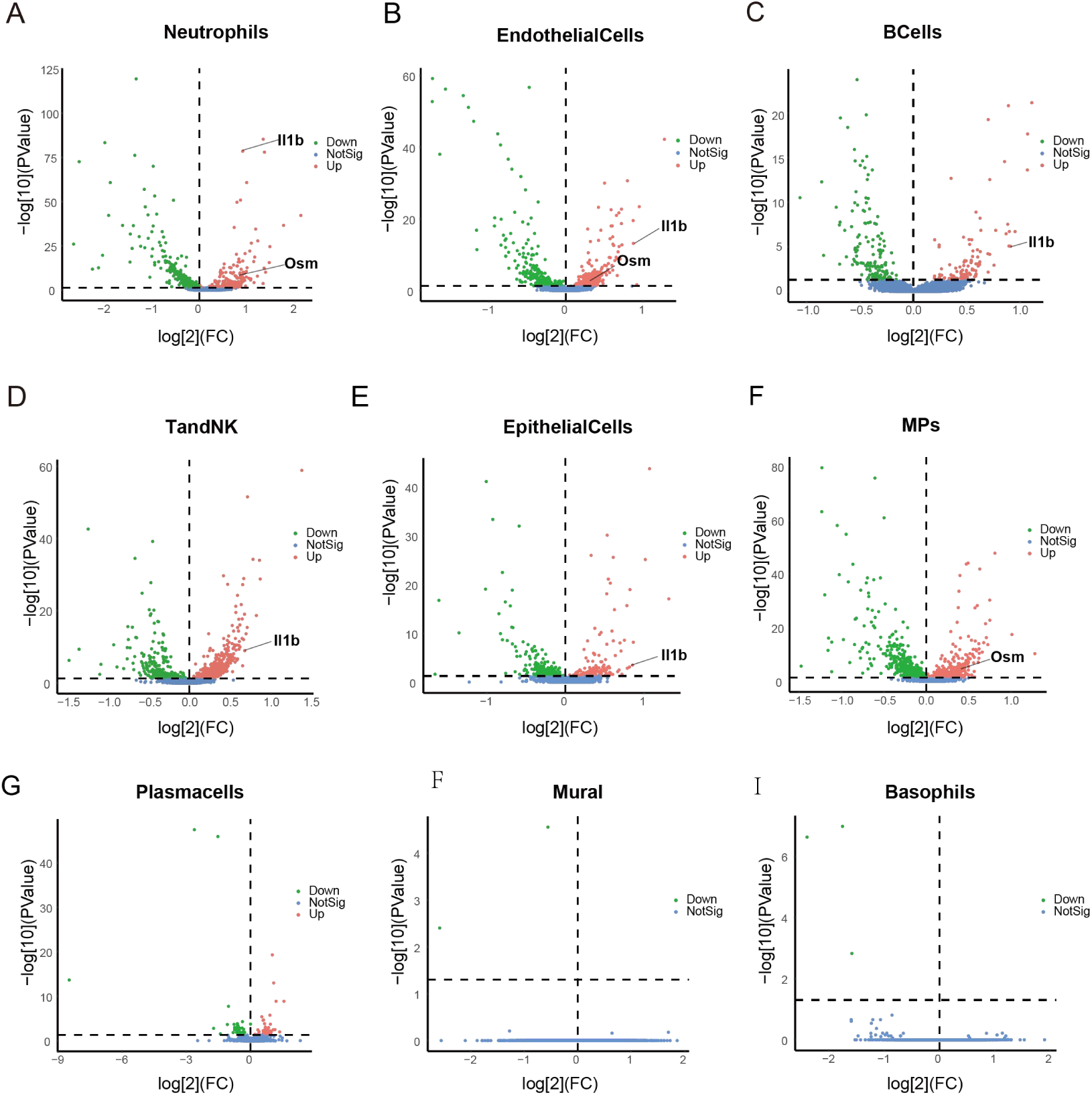
**Differential gene expression profiles of nine cell type.**(A) Volcano plot of differentially expressed genes in Neutrophils. (B)Volcano plot of differentially expressed genes in Endothelial cells.(C) Volcano plot of differentially expressed genes in B cells. (D)Volcano plot of differentially expressed genes in T and NK cells. (E) Volcano plot of differentially expressed genes in Epithelial cells.(F) Volcano plot of differentially expressed genes in Mononuclear phagocytes. (G) Volcano plot of differentially expressed genes in Plasma cells.(H) Volcano plot of differentially expressed genes in Mural cells.(I) Volcano plot of differentially expressed genes in Basophils.

Further analysis showed that the expression of IL1b was significantly up-regulated in Neutrophils, BCells, T/NK cells, EpithelialCells and endothelial cells. The expression of Osm was significantly up-regulated in neutrophils, endothelial cells and mononuclear phagocytes (MPS). Therefore, subsequent analyses targeting IL1b and Osm will focus on the cell types described above.

### 4. Expression of IL1b and Osm in Neutrophils and Mononuclear Phagocytes following PCV2d Infection and Their Potential Role in Pulmonary Fibrosis

This study focused on the expression patterns of key inflammatory factors IL1b and Osm in immune and stromal cells. IL1b expression was mainly detected in Neutrophils, B cells, natural killer cells, endothelial cells and epithelial cells. Osm expression analysis focused on neutrophils, endothelial cells (ECs) and myeloid phagocytic system cells (MPS cells).

High resolution single-cell transcriptome sequencing and visualization analysis showed that IL1b transcript abundance was at basal level or very low in B cells (S1 and S5 Figs), NK cells (S2 and S6 Figs), EP cells (S3 and S7 Figs) and ECs (S4 and S8 Figs), and the expression signal was significantly lower than the detection threshold, suggesting that these cells were not the main source of IL1b. Osm expression was significantly lower in ECs (S1 and S5 Figs).

Taken together, our preliminary results suggest that the core inflammatory signals of IL1b and Osm are mainly derived from neutrophils and MPS. The high expression of these two factors in the above cells suggests that they may play a dominant role in myeloid cell-mediated inflammatory response. As the key effector cells of the early immune response, neutrophils highly express these two factors at the same time, which deserves further study. The low contribution of B cells, NK cells, EP cells and ECs to IL1b expression, and the low expression of ECs and MPS cells to Osm further verified the localization of core cell groups.

As the key effector cells of the early inflammatory response, the inflammatory factors secreted by neutrophils may participate in the process of pulmonary fibrosis by regulating the activation of fibroblasts(13), and form a synergistic effect with the fibrosis mechanism driven by alveolar macrophage specific Osm. Single-cell transcriptome sequencing results showed that PCV2d infection significantly remodels the gene expression profile of neutrophils in lung tissue, with an overall upward trend of inflammation-related genes. Among them, the expression levels of IL1b and Osm, which are related to fibrosis, were significantly increased (Figs 4A and 4C). T-SNE visualization analysis confirmed a significant increase in the proportion of IL1b and Osm positive cells in the infected group (Fig 4B). Neutrophil subset analysis identified two functional subsets (Fig 4D) : Neutrophils_Ptgs2 and Neutrophils_Rpl12. Among them, the up-regulation of IL1b and Osm expression was mainly concentrated in the Neutrophils_Ptgs2 subset (Figs 4E and 4F), which also highly expressed inflammation-related genes (Fig 4G). The results showed that PCV2d infection specifically activated the Ptgs2 subset of neutrophils, which significantly upregulated the expression of IL1b and Osm. As a major subset accounting for more than 95%, Ptgs2 neutrophils are the core source of IL1b in the inflammatory microenvironment of pulmonary fibrosis, but their contribution to Osm needs to be comprehensively evaluated in combination with the subsequent analysis of mononuclear phagocytes.

**Figure 4.**
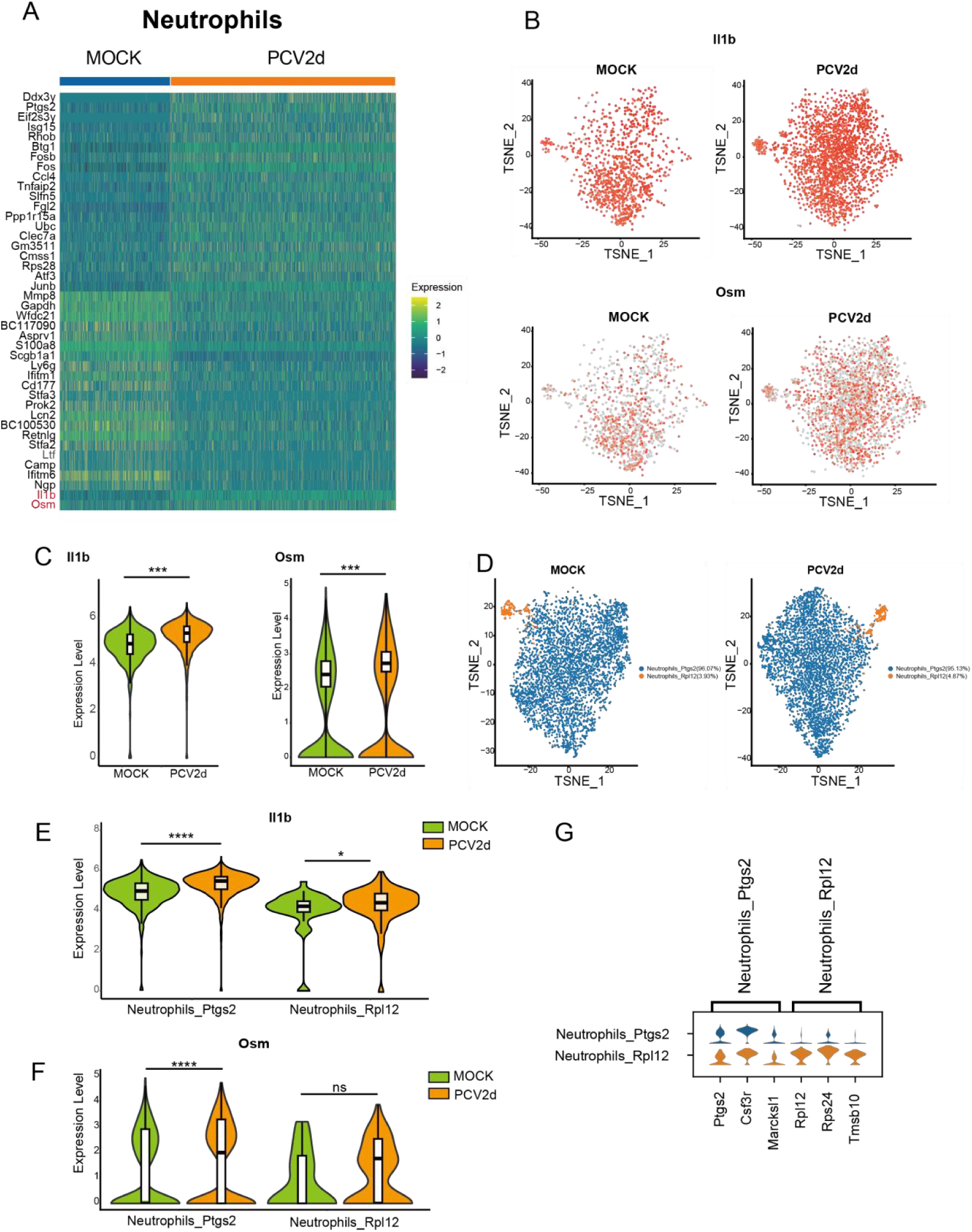
Analysis of IL1B and OSM expression levels in neutrophils from infected and healthy groups. (A) Neutro total differential genes under two treatments. (B) Expression of IL-1b and OSM in Neutro group of two treatment groups. (C) Violin expression of IL-1b and OSM genes in Neutro large classes under two treatment groups. (D) Neutro was grouped twice under two treatment groups. (E) Extraction of violin expression of IL1B in two Neutro subgroups (F)Extraction of violin expression ofOSM in two Neutro subgroups (G) Marker gene in two Neutro subgroups.

Mononuclear phagocytes (MPs) are an important population of immunomodulatory cells in the lung. Alveolar macrophages, a subset of MPS, are the main source of Osm and may play a central role in mediating the activation of fibroblasts and pulmonary fibrosis. scRNA-seq analysis revealed that PCV2d infection significantly upregulated the expression of Osm in mononuclear phagocytes (MPS) (Fig 5A). The transcript level of Osm in the infected group was significantly higher than that in the MOCK group (Fig 5C). T-SNE visualization analysis confirmed an increase in the proportion of Osm-positive cells (Fig 5B). MPS subgroup analysis identified five functional subsets (Fig 5D) : alveolar Macrophages, macrophages, Monocytes, cDC1, and cDC2. Among them, the AlveolarMacro subgroup showed the most significant upregulation of Osm expression in the infected group (Fig 5E), while other subsets, such as monocytes, showed no significant change in Osm expression. These results indicate that PCV2d infection can specifically induce Osm expression in alveolar macrophages, which is the main source of Osm in mononuclear phagocyte system, and directly supports the core conclusion that alveolar macrophage-specific Osm drives pulmonary fibrosis.

**Figure 5.**
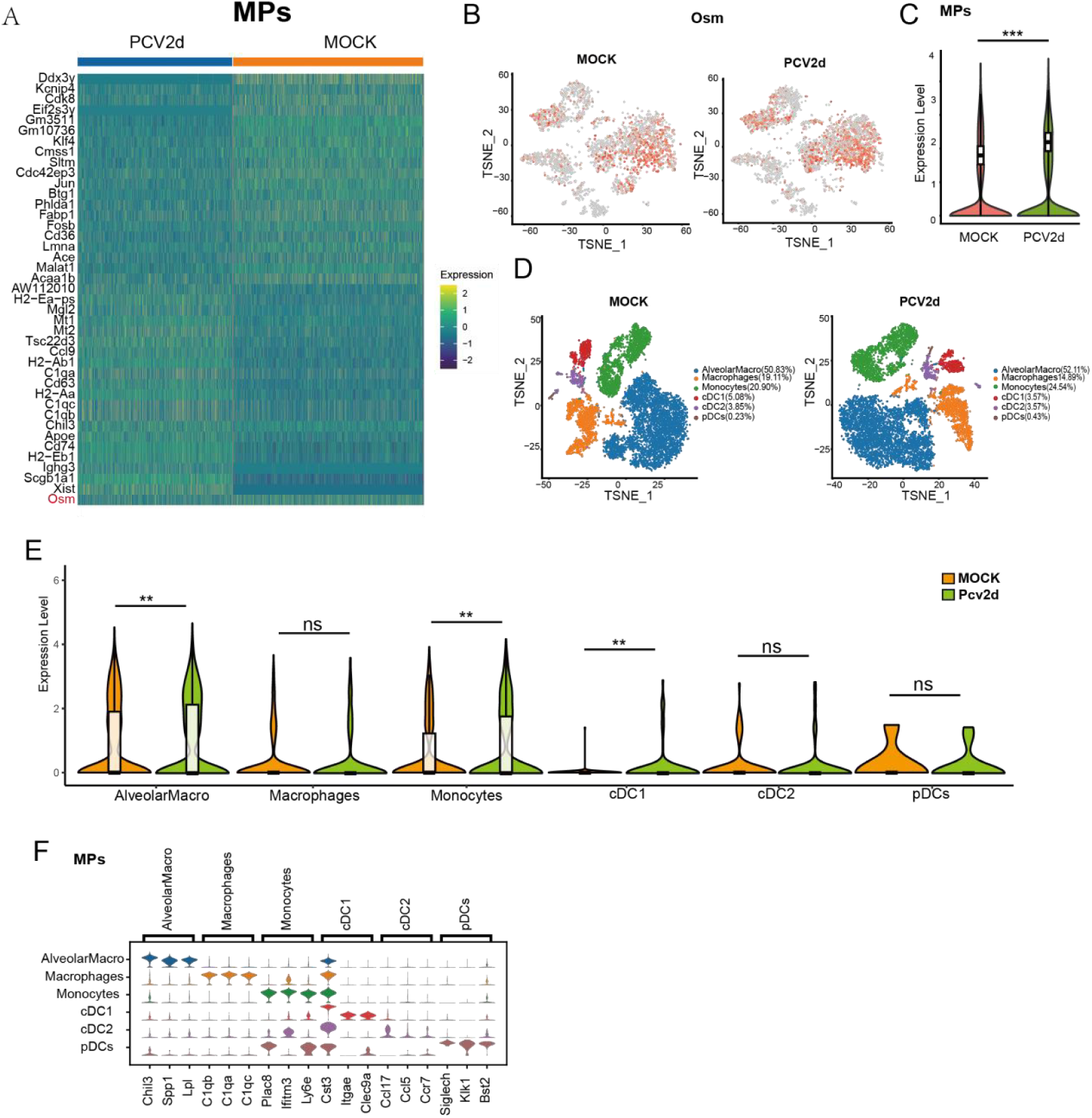
Analysis of OSM expression level in mononuclear phagocytes of infected and healthy groups. (A) MPS total difference genes under two treatments. (B) OSM gene violin expression of MPS large class under two treatment groups. (C) Violin expression of OSM gene in MPS class. (D) Secondary grouping of MPS under two treatment groups. (E) Extraction of violin expression quantity of OSM in MPS two subgroups. (F) Marker genes for both subgroups.

### 5. Histopathological Phenotype Analysis of Pulmonary Fibrosis after Vaccine Intervention

To assess the protective effect of vaccine intervention on PCV2d induced pulmonary fibrosis and its effect on alveolar macrophage expression of Osm and fibroblast activation, control experiments were established using BALB/c mice. As shown in Fig 6A, mice were divided into two groups: adjuvant control group (Adjuvant group) and vaccine treatment group (Vaccine group). Histopathological results (Fig 6B) showed significant differences between the two groups: the vaccine-treated group had intact lung tissue structure, clear alveoli and no significant interstitial thickening, while the adjuvant control group showed typical fibrosis features, including alveolar collapse, interstitial widening and inflammatory cell infiltration. ImageJ quantitative analysis (Fig 6C) confirmed that the alveolar vacuolation area of the vaccine-treated group was significantly larger than that of the adjuvant control group, indicating that the vaccine was effective in reducing PCV2d induced lung parenchyma damage and fibrosis.

**Figure 6.**
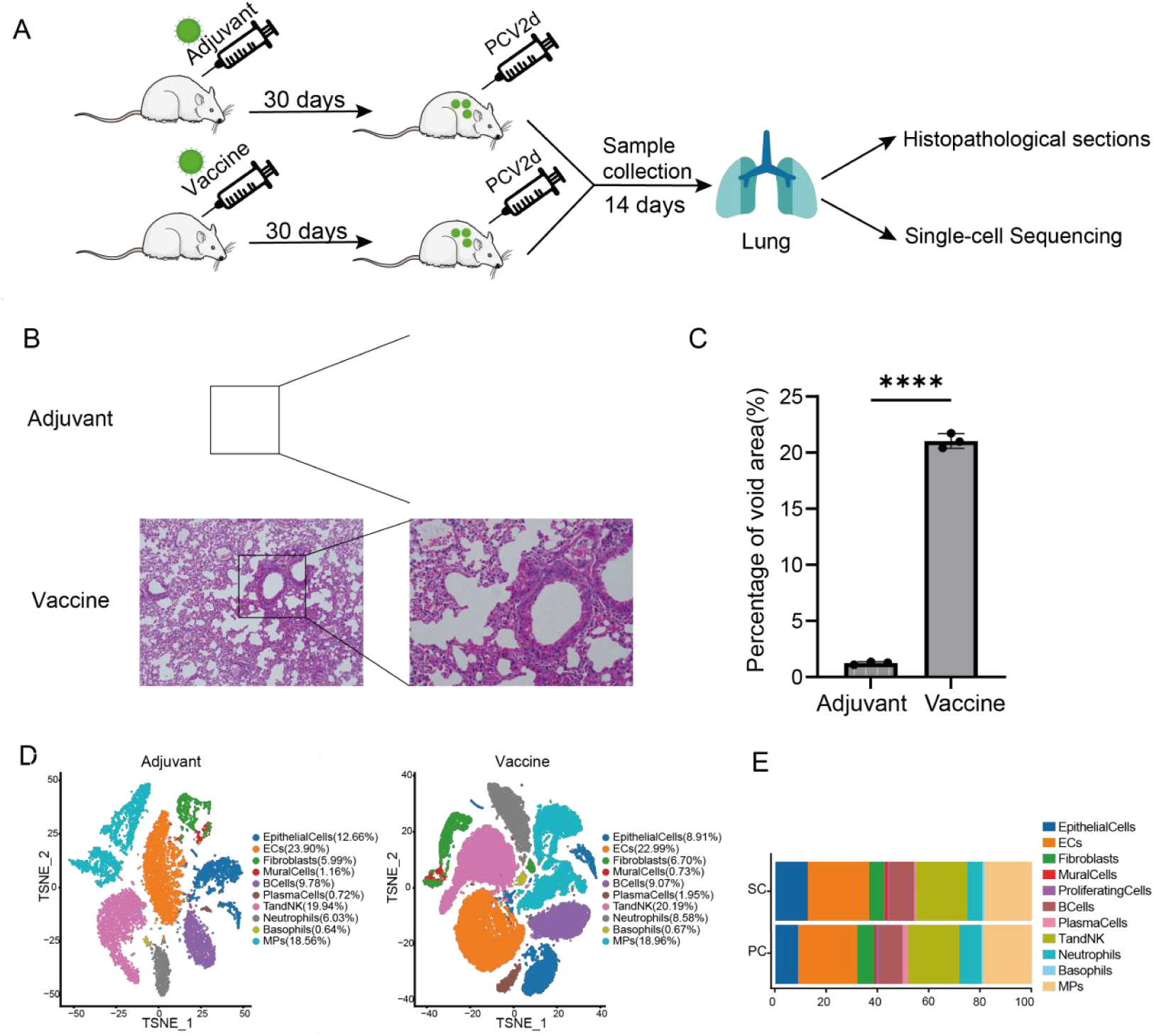
Design and differences in lung cell composition investigated throughout vaccine intervention in BALB/c mice. (A) Test flow design. (B) Differences in lung fibrosis sections between adjuvant group and vaccine group. (C) Count the percentage of vacuoles in three random visual fields. (D) Cell grouping in both treatment groups. (E) Percentage of each subgroup in both treatment groups.

Single cell transcriptome analysis (Fig 6D) further revealed changes in lung tissue cell populations. Cell proportion statistics (Fig 6E) showed a significantly higher proportion of neutrophils and B cells in the vaccine-treated group, suggesting an enhanced acute immune response, while the proportion of fibrotic core effector cells fibroblasts decreased in the vaccine-treated group, consistent with a reduction in fibrotic phenotype. These changes in cell populations provide a preliminary basis for further verification of whether the vaccine affects fibroblast activation by regulating Osm signaling pathway in alveolar macrophages. The results showed that PCV2d vaccine intervention could significantly alleviate the degree of pulmonary fibrosis in challenged mice, manifested by preservation of lung tissue structure, increase of alveolar vacuolation area and decrease of fibroblast proportion. Single cell analysis showed dynamic changes in immune cell populations, increased neutrophils and B cells, and decreased fibroblasts, laying a foundation for further investigation into whether the vaccine plays a protective role by inhibiting Osm signaling in alveolar macrophages.

### 6. Analysis of vaccine intervention on expression of Col1a2 and Timp3 in fibroblasts

To explore the effect of Vaccine administration after PCV2d infection on lung fibroblast function, gene expression profiles of adjuvant control group (Adjuvant) and vaccine treatment group (Vaccine) were systematically compared by single cell transcriptome sequencing technology (Fig 7A). Heat map analysis showed that the expression levels of Col1a2 and Timp3 decreased significantly after PCV2d infection.

**Figure 7.**
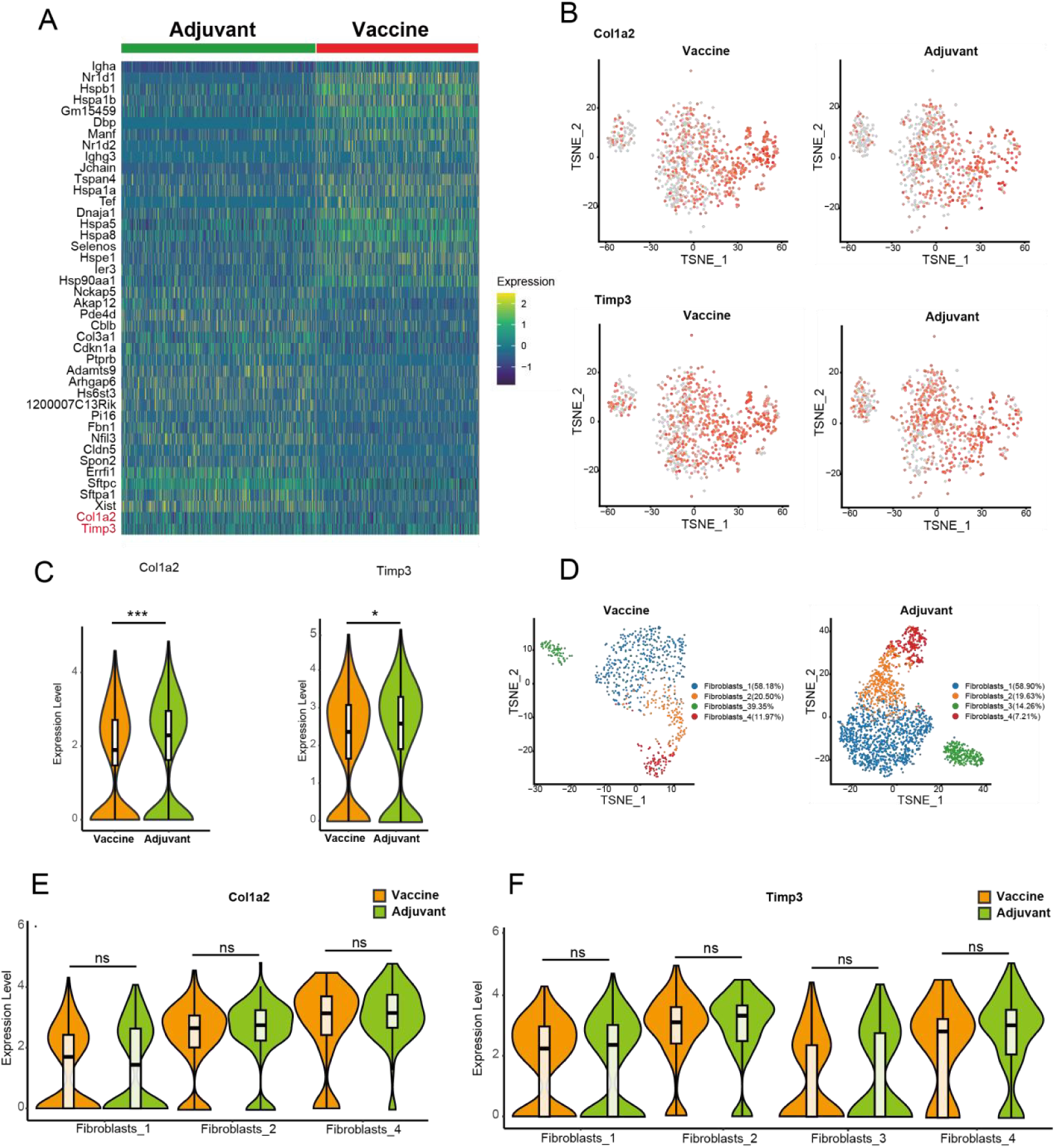
Analysis of fibrosis factors in Adjuvant and Vaccine groups in pulmonary fibrosis phenotypes under vaccine intervention. (A) Total differential genes in fibroblasts under two treatments. (B) Fibroblast factors COL1A2 and TIMP3 were expressed in the fibroblast population of both treatment groups. (C) Violin expression levels of COL1A2 and TIMP3 genes in fibroblasts under both treatment groups. (D) Secondary grouping of fibroblasts under both treatment groups. (E) Marker genes of four subgroups. (F) Violin expression of COL1A2 in each subgroup under both treatment groups. (G) TIMP3 Violin expression in each subgroup under both treatment groups.

Visual analysis using TSNE dimensionality reduction (Fig 7B) showed that the proportion of Col1a2 and Timp3 positive cells was significantly reduced in the Vaccine group compared with the Adjuvant group. Further quantitative analysis of violin plots (Fig 7C) confirmed that the expression intensity of these two marker fibrosis-related genes was significantly reduced in the Vaccine group compared with the Adjuvant group.

In-depth analysis of the changes in fibroblast subsets revealed that vaccination after PCV2d infection significantly changed the composition of cell subsets (Fig 7D).Among them, 11.97% of fibroblasts4 and 39.35% and 14.26% of fibroblasts3 were in the vaccine and adjuvant groups respectively.

Violin plot quantitative analysis of different subsets of fibroblasts (Figs 7E and 7F) demonstrates differences in gene expression among different subsets of cells after vaccination. There was no significant difference in the expression of Col1a2 and Timp3 in different neutrophil subsets, indicating that the contribution of each fibroblast subset to Col1a2 and Timp3 was the same in Adjuvant and Vaccine groups.

### 7. Vaccine Intervention Does Not Alter IL1b and Osm Expression in Neutrophils but Specifically Suppresses Osm in Alveolar Macrophages

To explore the effect of vaccination on neutrophils in lung tissue, we analyzed the gene expression profile of neutrophils in lung tissue by single-cell RNA sequencing. The results showed that vaccination significantly altered the gene-expression profile of neutrophils (Fig 8A). T- SNE dimensionality reduction visualization (Fig 8B) showed no obvious change in the clustering distribution of cells between the Vaccine group and the Adjuvant group, and there was also no significant difference in the proportion of IL1b - positive and Osm - positive cells. Violin plots (Fig 8C) were used to display the expression levels of IL1b and Osm in the two groups, and no significant differences were observed. Further sub - population analysis (Fig 8D) indicated that the proportions of specific sub - populations such as Neutrophils_Ptgs2 and Neutrophils_Rpl12 did not change significantly. Quantitative analysis of violin plots for different neutrophil sub - populations (Fig 8E) demonstrated the gene expression differences of neutrophils after infection, yet no obvious differences were found in the expression of IL1b and Osm among different sub - populations. Overall, multi - dimensional analyses revealed the significant impact of vaccination on the gene expression profiles and sub - population composition of macrophages.

**Figure 8.**
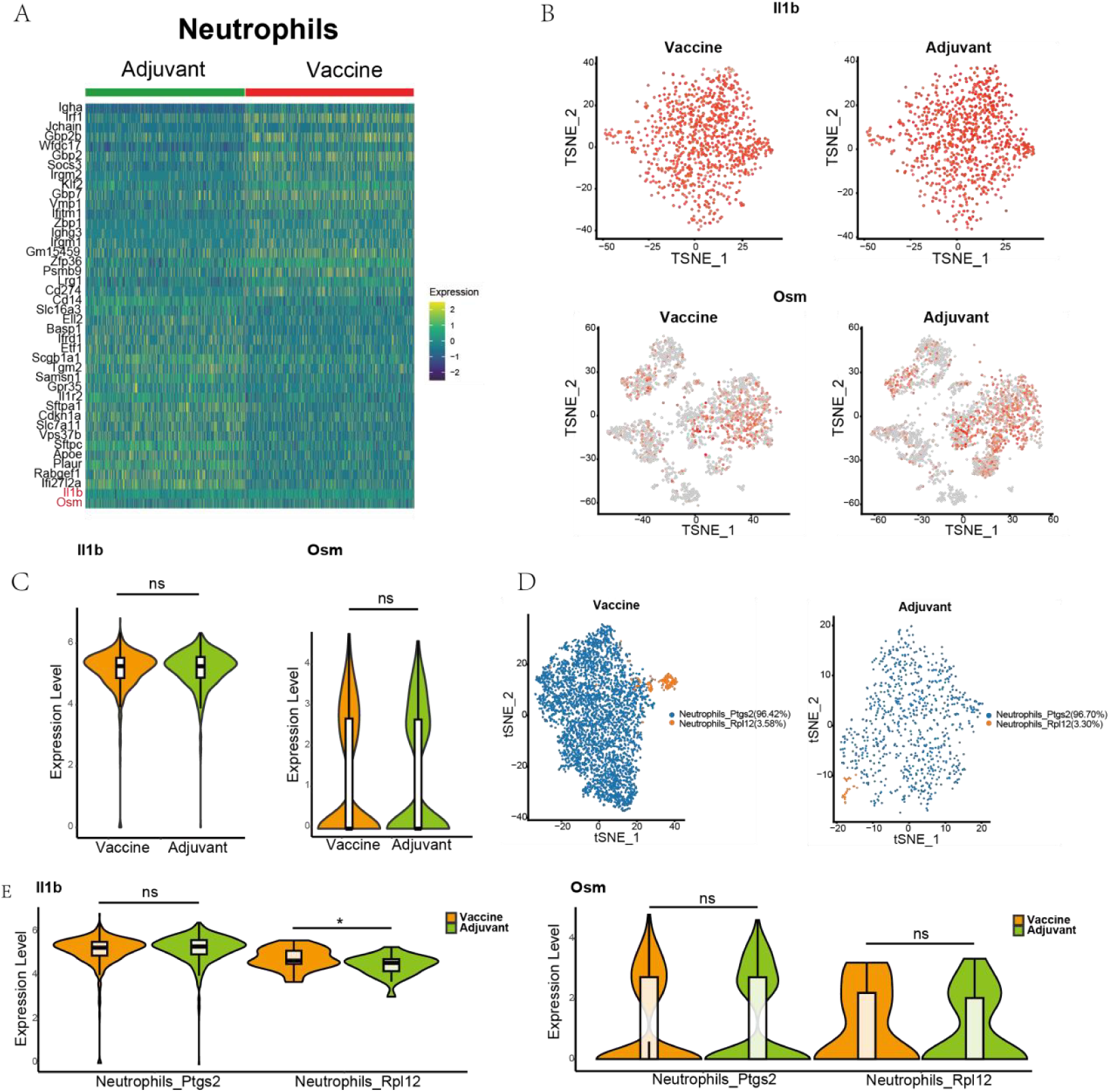
Analysis of IL1B and OSM expression levels in Neutrophils of Adjuvant and Vaccine groups. (A) Neutro total differential genes under two treatments. (B) Expression of IL-1b and OSM in Neutro group of two treatment groups. (C) Violin expression of IL-1b and OSM genes in Neutro large classes under two treatment groups. (D) Neutro was grouped twice under two treatment groups. (E) Extraction of violin expression of IL1B and OSM in Neutro subgroups.

In contrast, vaccination induced significant changes in the transcriptional landscape and subpopulation composition of macrophages. A gene expression heatmap clearly depicted distinct transcriptional profiles between the Vaccine and Adjuvant groups (Fig 9A), including a significant reduction in Osm expression within the Vaccine group.

**Figure 9.**
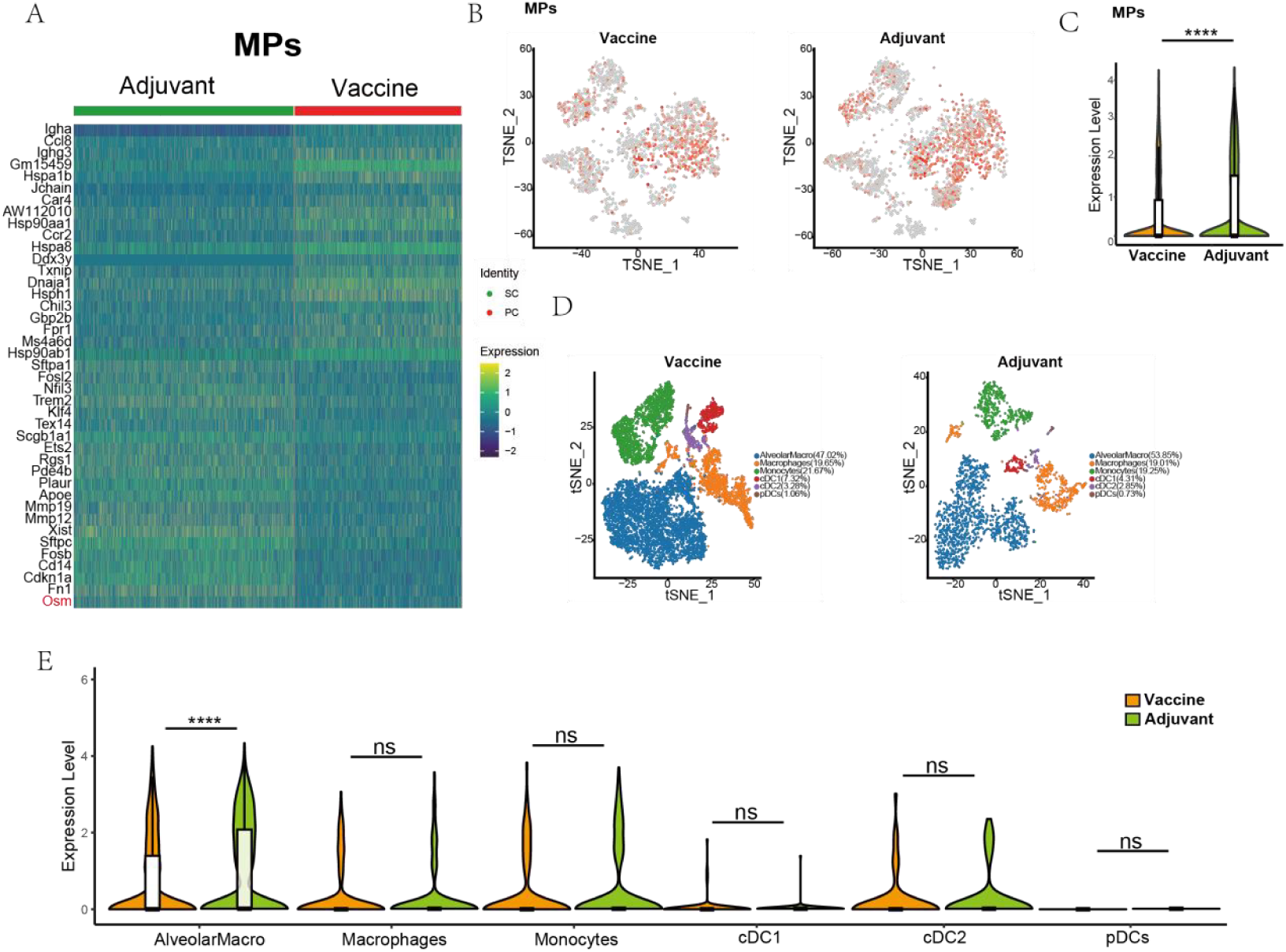
Analysis of OSM expression level in mononuclear phagocytes of Adjuvant and Vaccine groups. (A) MPS total difference genes under two treatments. (B) OSM expression in MPS group of two treatment groups. (C) OSM gene violin expression of MPS large class under two treatment groups. (D) Secondary grouping of MPS under **two** treatment groups. (E) Violin expression of OSM in MPS subgroups.

T-SNE-based dimensionality reduction and clustering analysis confirmed a marked shift in cellular clustering distribution for macrophages in the Vaccine group compared to the Adjuvant group (Fig 9B). This shift was accompanied by a significant decrease in the proportion of Osm-positive cells. Violin plots of differential gene expression demonstrated that Osm levels were significantly lower in the Vaccine group versus the Adjuvant group (Fig 9C).

Analysis of macrophage subsets by T-SNE visualization revealed an altered distribution pattern in the vaccine group (Fig 9D), with significant changes in the proportions of key subsets, including alveolar Macrophages, macrophages, and Monocytes. Violin plot analysis of Osm expression in each macrophage subset showed that the alveolar macrophage subset showed higher Osm expression as a whole, while the other subsets showed similar expression levels of Osm in the adjuvant control and vaccine groups. This suggests that vaccine-mediated inhibition of Osm is alveolar macrophage subpopulation specific.

### 8. In Vitro Co-Culture Model Validates the Fibrotic Regulatory Mechanism

To verify the feasibility of the key regulatory mechanisms of pulmonary fibrosis in a cross-species model, a porcine macrophage-lung fibroblast heterologous co-culture system (Fig 10E) was constructed to simulate the microenvironment interaction scenario of viral infection. Using single-cell sequencing, we identified Osmr, the receptor for Osm, on PLF (Figs 10A-D), and then performed the following experiments: 24 hours after the co-culture system was inoculated with PCV2d virus, Viral load (PCV2d) and expression dynamics of fibrosis-related molecular markers (*COL1A2, TIMP3, IL1b, OSM*) were detected by qRT-PCR system (Figs 10G-I). Among them, PCV2d detection was used to confirm the efficiency of virus infection, and the results showed that PCV2d increased significantly after infection, confirming the successful establishment of the model.

**Figure 10.**
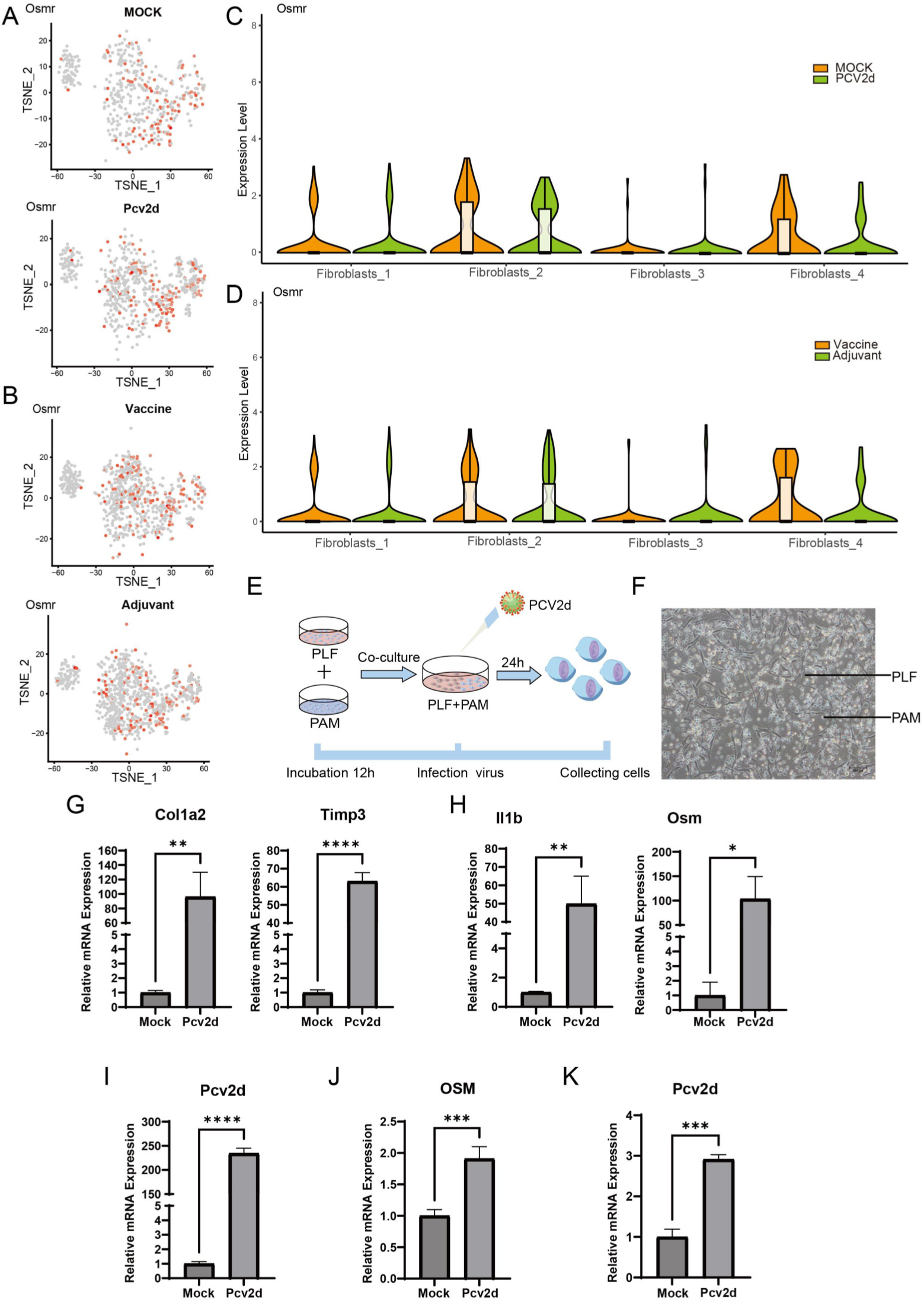
In vitro co-culture model to validate the regulatory mechanism of fibrosis. (A) Expression of Osmr in the large categories of fibroblasts in the MOCK group, Pcv2d group, Vaccine group, and Adjuvant group. (B) Osmr expression in the large categories of fibroblasts inthe Vaccine group and the Adjuvant group. (C) Violin expression of Osmr gene in MPS major classes in MOCK and Pcv2d groups. (D) Violin expression of Osmr in MPS subgroup under MOCK and Pcv2d groups. (E) Schematic representation of PAM and PLF co-culture. (F) Diagram of co-cultured cells. (G) Gene expression of Col1a2 and Timp3 in co-cultured cells infected with PCV2d. (H) IL1b and Osm gene expression in co-cultured cells infected with PCV2d. (I) Detection of viral expression in co-cultured cells infected with PCV2d. (J) Osm gene expression after PAM infection of PCV2d. (K) Detection of viral expression after PAM infection of PCV2d.

Quantitative analysis showed that PCV2d infection significantly activated molecular networks of fibrosis-related pathways (Figs 10G and 10H). Specifically, the expression of *COL1A2* and *TIMP3* increased significantly. Meanwhile, *IL1b* and *OSM*, the key mediators of inflammatory microenvironment, also showed a significant increase trend.

To determine whether *OSM* is secreted by PAM or PLF, PAM and PLF were cultured separately and treated with PCV-2 virus. After 24 hours, qRT-PCR analyses of viral load and *OSM* expression (Figs 10J and 10K) demonstrated that *OSM* was predominantly expressed in PAMs, with negligible expression detected in PLFs, confirming macrophages as the primary cellular source of *OSM* in this context.

## Discussion

Our study employs single-cell RNA sequencing (scRNA-seq), histopathology, and in vitro co-culture models to systematically dissect the cellular and molecular mechanisms underlying PCV2d-induced pulmonary fibrosis (PF). We identified alveolar macrophage (AlveolarMacro)-specific oncostatin M (Osm) as a key driver of fibroblast activation and extracellular matrix (ECM) deposition, with vaccine intervention targeting this pathway significantly alleviating fibrotic progression (Fig 11). These findings provide novel insights into viral-induced pulmonary fibrosis (PF) and potential therapeutic strategies.

**Fig 11.**
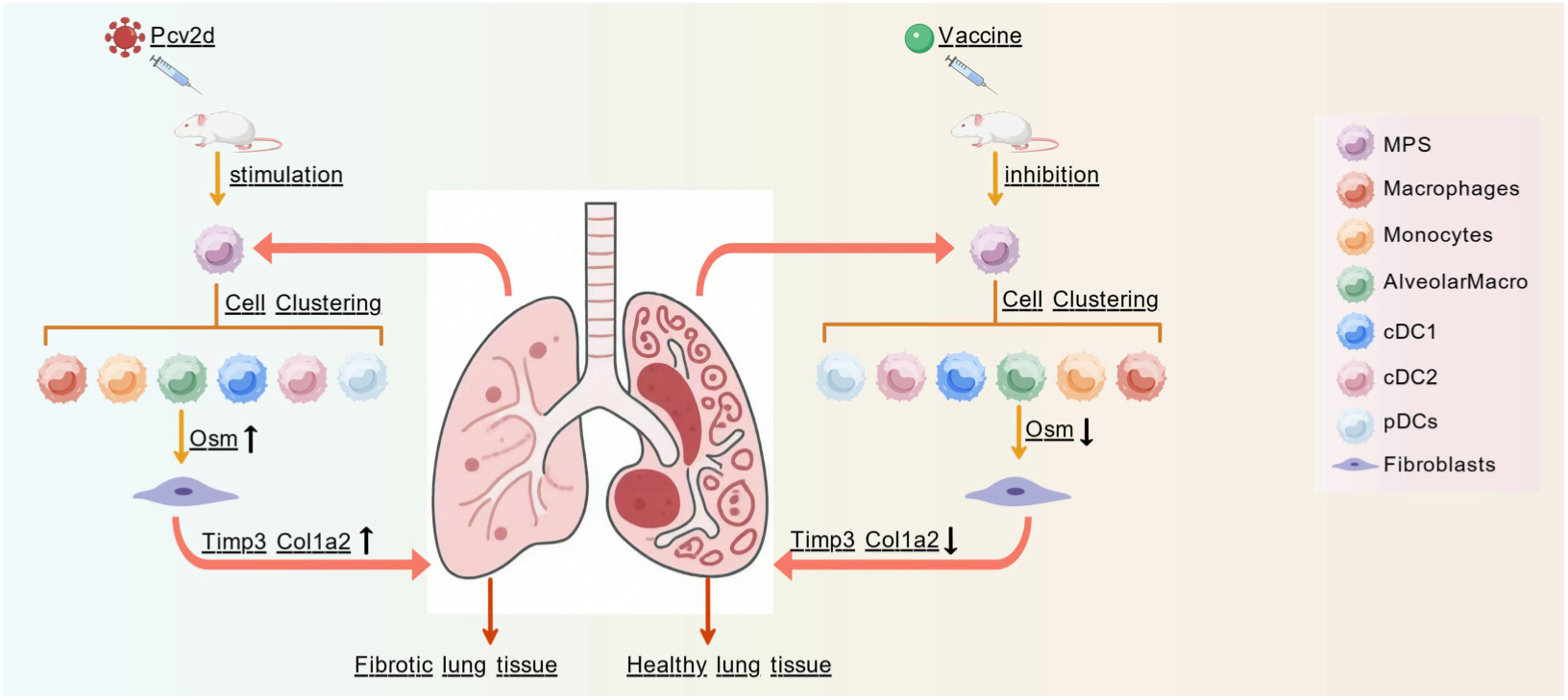
**PCV2d infection affects pulmonary fibrosis.**

### 1. Fibroblast activation as a central event in PCV2d-induced PF

Fibroblasts are pivotal effectors in PF, with their pathological activation leading to excessive ECM production. Our scRNA-seq analysis revealed that PCV2d infection specifically upregulates the expression of Col1a2 (encoding type I collagen) and Timp3 (a matrix metalloproteinase inhibitor) in lung fibroblasts, consistent with enhanced ECM synthesis and remodeling. Subcluster analysis further identified Fibroblasts_1 as the key pro-fibrotic subset, characterized by co-expression of Col1a2, Timp3, and the myofibroblast marker Acta2. This subset was significantly expanded in infected lungs, confirming its role as a central mediator of ECM deposition.

These observations align with prior studies demonstrating that viral infections can trigger fibroblast activation through paracrine signaling(14). Importantly, our data extend this knowledge by linking PCV2d infection to a specific fibroblast subpopulation, highlighting Fibroblasts_1 as a potential therapeutic target for inhibiting ECM accumulation.

### 2. Alveolar macrophage-specific **Osm**: A critical pro-fibrotic cytokine

Inflammatory cytokines are well-recognized regulators of fibroblast function in PF(11). Through multi-omics screening, we identified Osm and IL1b as core pro-fibrotic factors in PCV2d-infected lungs. While IL1b was primarily derived from Neutrophils_Ptgs2, Osm expression was selectively enriched in AlveolarMacro within the mononuclear phagocyte system (MPS).

Notably, scRNA-seq trajectory analysis and subcluster profiling revealed that PCV2d infection specifically upregulates Osm in AlveolarMacro, whereas other MPS subsets (e.g., monocytes, cDC1) showed no significant changes. Functional validation via in vitro co-culture models further confirmed that Osm secreted by porcine alveolar macrophages (PAMs) – the porcine homolog of AlveolarMacro – directly drives pulmonary fibroblast (PLF) activation, as evidenced by increased Col1a2 and Timp3 expression. These results establish AlveolarMacro-derived Osm as a non-redundant driver of PCV2d-induced fibroblast activation.

### 3. Vaccine intervention alleviates PF by targeting AlveolarMacro Osm

Vaccination is a proven strategy to mitigate viral pathogenesis, but its impact on viral-induced fibrosis remains understudied. Our vaccine intervention experiments demonstrated that PCV2d inactivated vaccine significantly preserved lung architecture, reduced interstitial thickening, and increased alveolar vacuolization – hallmarks of reduced fibrotic severity. Single-cell analysis linked these phenotypic improvements to a decreased proportion of pro-fibrotic fibroblasts and, critically, suppressed Osm expression in AlveolarMacro.

Notably, vaccine intervention did not alter IL1b or Osm levels in neutrophils, suggesting its specificity for the AlveolarMacro-Osm axis. This selective inhibition aligns with the observed reduction in Fibroblasts_1 and ECM markers (Col1a2, Timp3), reinforcing that targeting AlveolarMacro Osm is sufficient to blunt fibrotic progression. These data provide preclinical evidence that vaccines can modulate the fibrotic microenvironment beyond viral clearance, offering a dual benefit in viral PF.

### 4. Cross-species validation supports the conserved role of Osm

To strengthen the translational relevance of our findings, we established a porcine PAM-PLF co-culture model, which recapitulated key features of PCV2d-induced fibrosis. PCV2d infection in this system upregulated *OSM* in PAMs (but not PLFs) and induced fibroblast activation, mirroring our in vivo observations in mice. This cross-species consistency suggests that the AlveolarMacro-*OSM*-fibroblast axis is evolutionarily conserved, supporting its potential relevance to human viral PF (e.g., SARS-CoV-2 or influenza-induced fibrosis).

### Limitations and future directions

Our study has limitations. First, we focused on Osm and IL1b, but other cytokines may contribute to the fibrotic microenvironment. Second, the long-term dynamics of AlveolarMacro Osm expression and its impact on fibrosis resolution require further investigation. Third, while our vaccine data support Osm as a target, direct Osm neutralization experiments would formally confirm causality.

Future studies could explore combinatorial strategies targeting Osm and IL1b to synergistically inhibit fibrosis, or investigate epigenetic regulation of Osm in AlveolarMacro during PCV2d infection. Additionally, translating these findings to human viral PF models (e.g., organoids or patient-derived cells) would enhance clinical relevance.

## Conclusion

In summary, our work identifies AlveolarMacro-specific Osm as a critical driver of PCV2d-induced pulmonary fibrosis, mediated through fibroblast activation. Vaccine intervention targeting this pathway alleviates fibrosis, highlighting Osm as a promising therapeutic target. These findings advance our understanding of viral-induced PF and lay the groundwork for developing precision therapies.

## Abbreviations

Acronym: Full Name
Col1a2: Encoding type I collagen
Timp3: Matrix metalloproteinase inhibitor
IL1b: Interleukin-1B
Osm: Oncostatin M
PCV2d: The porcine circovirus type 2d
PF: Pulmonary fibrosis
ECM: Extracellular matrix
MPS: Mononuclear phagocytes
ECs: Endothelial cells
NK cells: TandNK cells
EP cells: Epithelial Cells
PAM: Macrophage
PLF: Fibroblast
DMEM: Dulbecco’s Modified Eagle Medium

## Data availability statement

All relevant data can be found within this article and mice lungs single-cell RNA-seq data is obtained from the ScienceDB database (https://doi.org/10.57760/sciencedb.30497).

## ARRIVE guidelines

The manuscript has adhered to ARRIVE guidelines.

## Ethics statement

The animal experiment was approved by the Northwest A&F University Institutional Animal Care and Use Committee (approval number: XN2023-0903). The research was conducted in compliance with local laws and institutional regulations.

## Author contributions

JX: Data curation, Methodology, Formal analysis, Writing – original draft. XF: Data curation, Writing – original draft. PC: Data curation, Writing – original draft.YY: Formal analysis, Writing – original draft.YC: Formal analysis, Writing – original draft.GF: Formal analysis, Writing – original draft.BY: Formal analysis, Writing – original draft.FY: Formal analysis, Writing – original draft.YW: Formal analysis, Writing – review & editing. SZ: Methodology, Formal analysis, Writing – review & editing.

## Generative AI statement

The author(s) declare that no Gen AI was used in the creation of this manuscript.

## Acknowledgments

We would like to acknowledge the staff at Shaanxi Stem Cell Engineering Research Center for technical assistance and the supporting staff - Su Ang and Liu Pengfei at Singleron Biotechnologies for assistance in data collection.

---

**Supplementary Figure 1:**
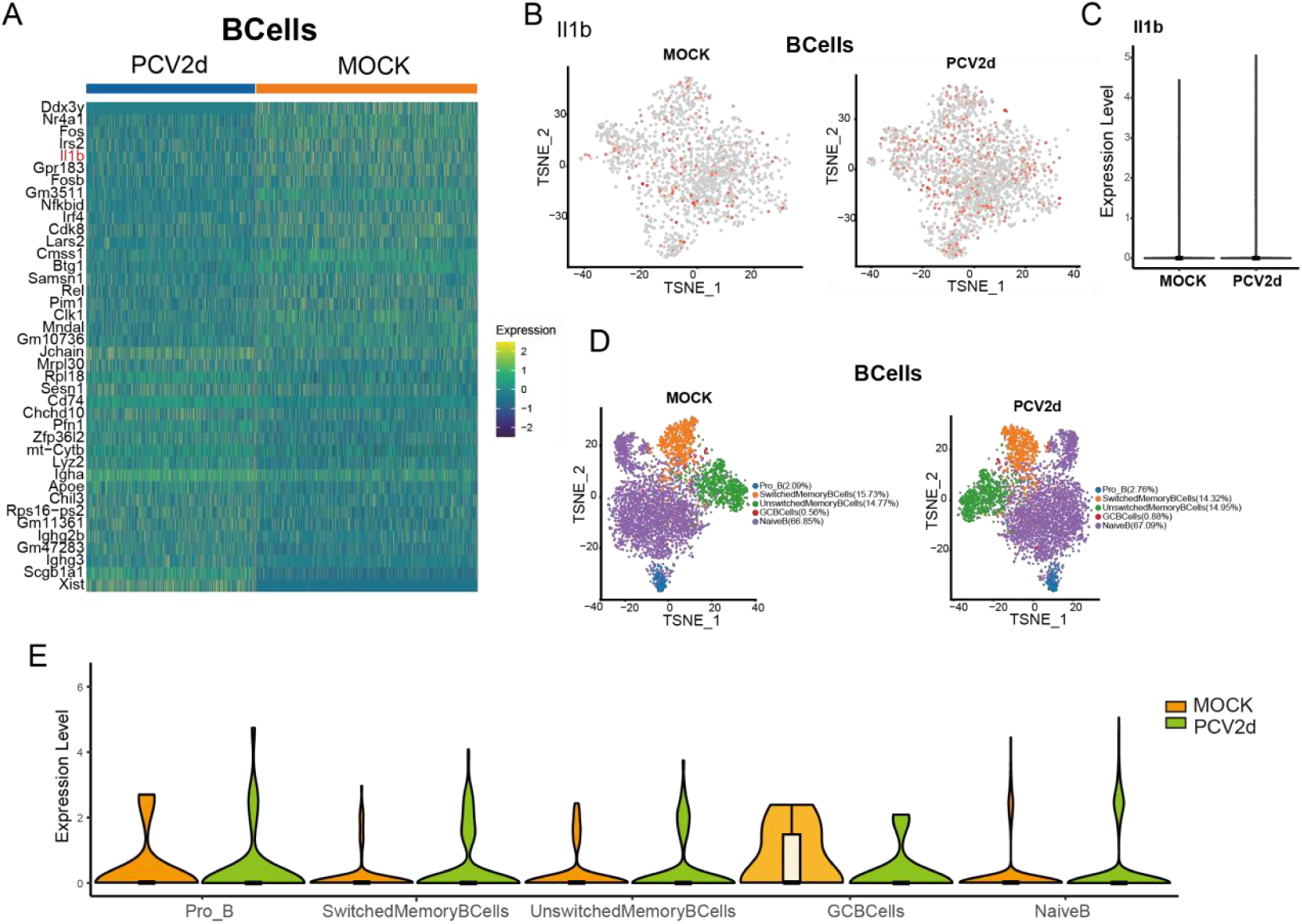
Analysis of B cells in MOCK group and PCV2d infection group. (A) Total differential genes in Bcells under two treatments. (B) IL-1b expression in B-cells of both treatment groups. (C) IL-1b gene violin expression of cello in Bcells under two treatment groups. (D) Secondary grouping of Bcells under two treatment groups. (E) Violin expression of IL-1b in two subsets of B cells.

**Supplementary Figure 2:**
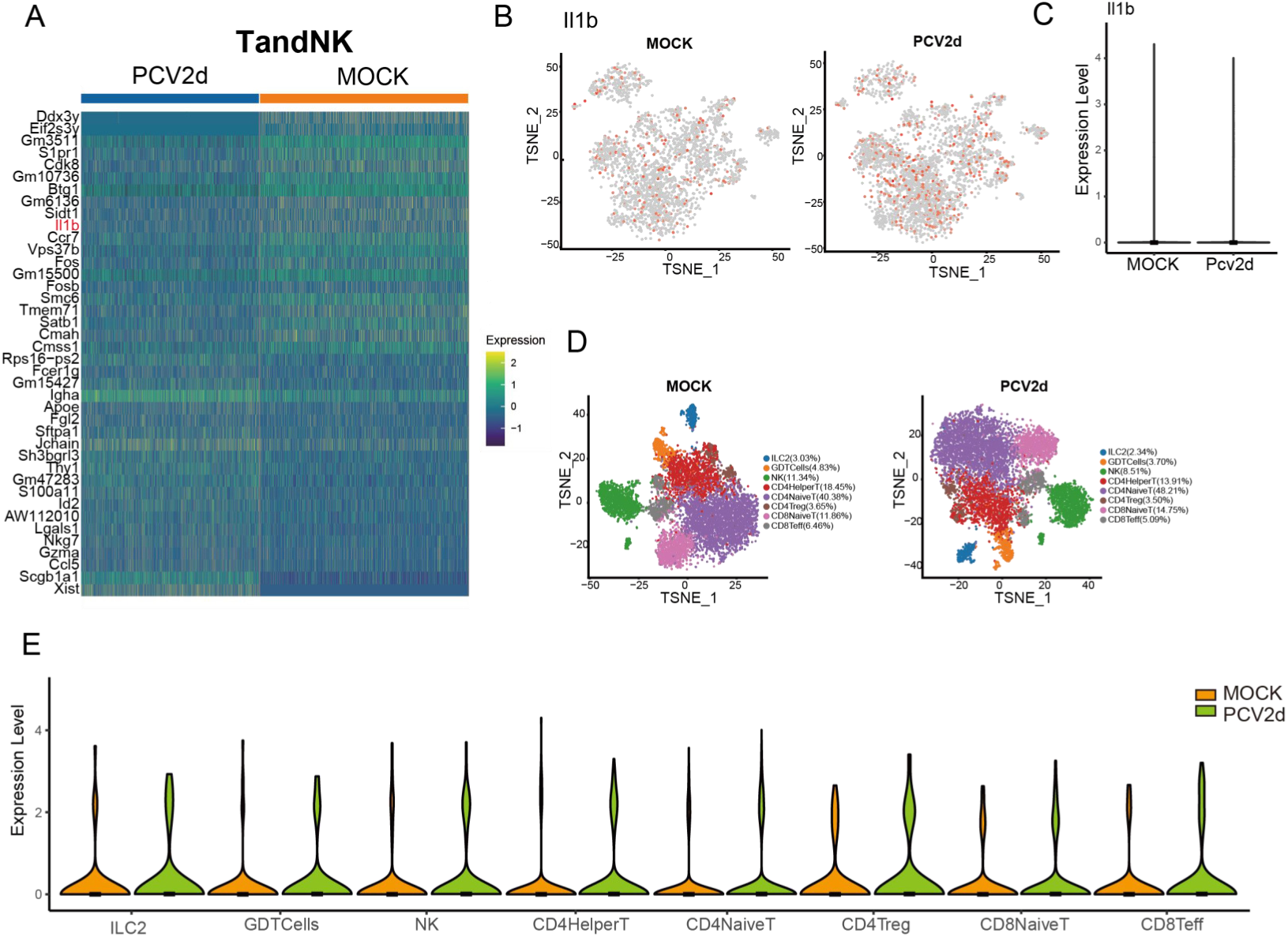
TandNK analysis of MOCK group and PCV2d infection group. (A) TandNK total differential genes under two treatments. (B) IL1b expression in TandNK class of two treatment groups. (C) Violin expression of IL1b gene of TandNK cello class under two treatment groups. (D) TandNK was divided into two groups. (E) Violin expression of IL1b in T and NK subpopulations.

**Supplementary Figure 3:**
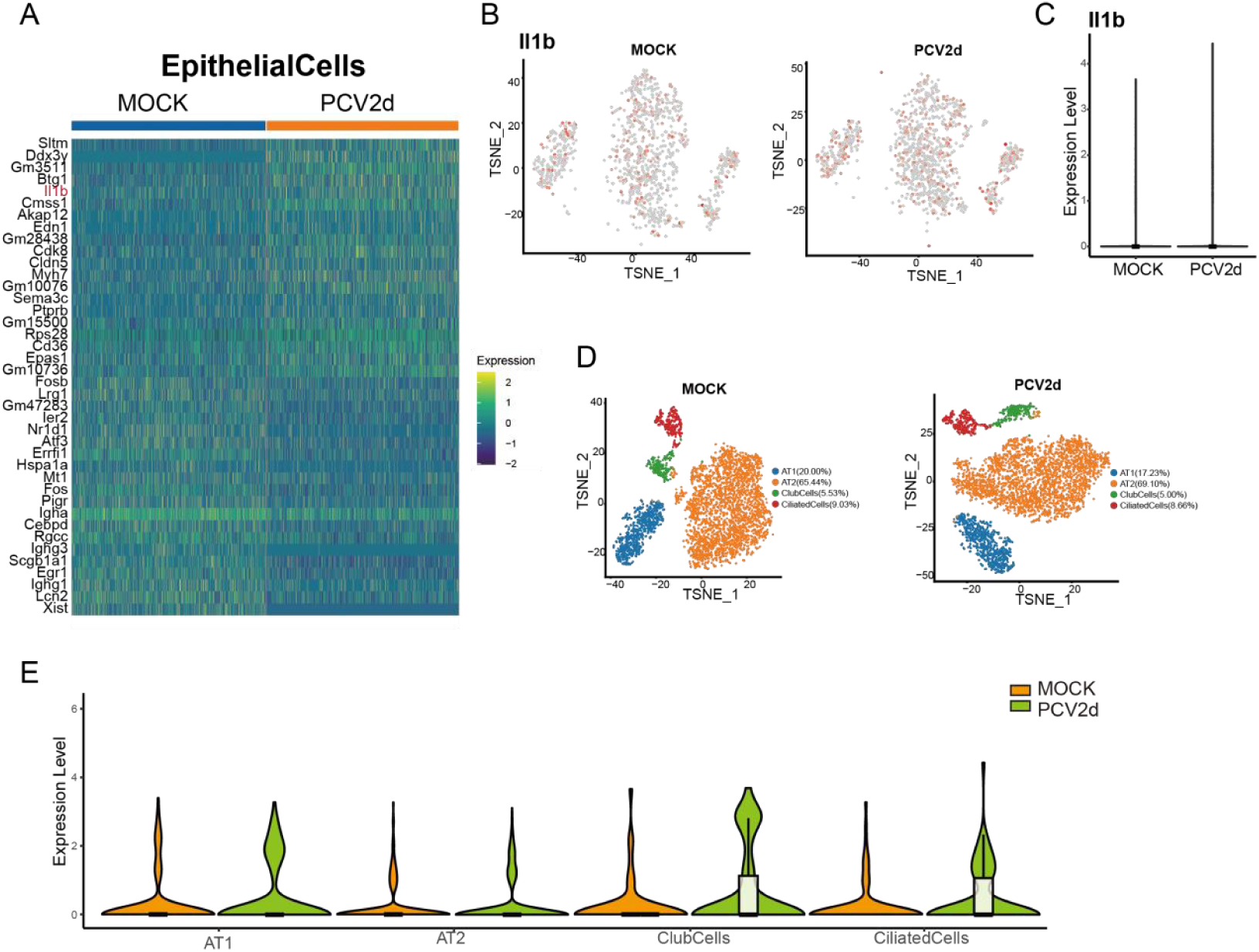
EpithelialCells analysis of MOCK group and PCV2d infection group. (A) Total differential genes in EpithelialCells under two treatments. (B) IL1b expression in EpithelialCells of both treatment groups. (C) IL1b gene violin expression in EpithelialCells under both treatment groups. (D) Subgrouping EpithelialCells under two treatment groups. (E) Violin expression of IL1b in two subgroups of EpithelialCells.

**Supplementary Figure 4:**
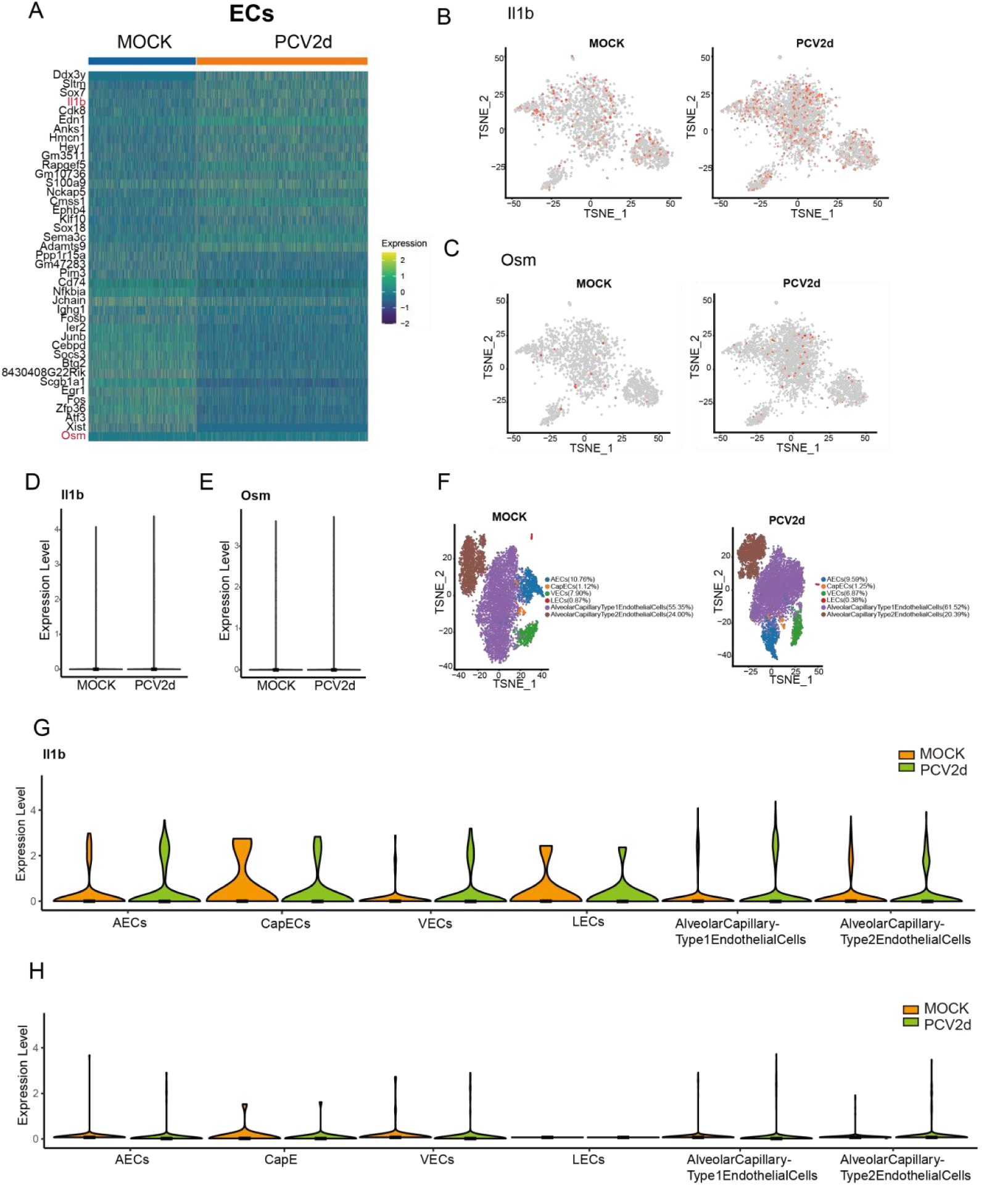
ECs analysis of the MOCK group and the PCV2d infection group. (A) Total differential genes for ECs under two treatments. (B) IL1b was expressed in the ECs category of the two treatment groups. (C) Osm was expressed in the ECs category of the two treatment groups. (D) Violin expression of IL1b gene in ECs under the two treatment groups. (E) Violin expression of Osm gene in ECs under the two treatment groups. (F) ECs were sub-grouped under the two treatment groups. (G) Violin expression of IL1b in two subpopulations of ECs, and (H) Violin expression of Osm in two subpopulations of ECs.

**Supplementary Figure 5:**
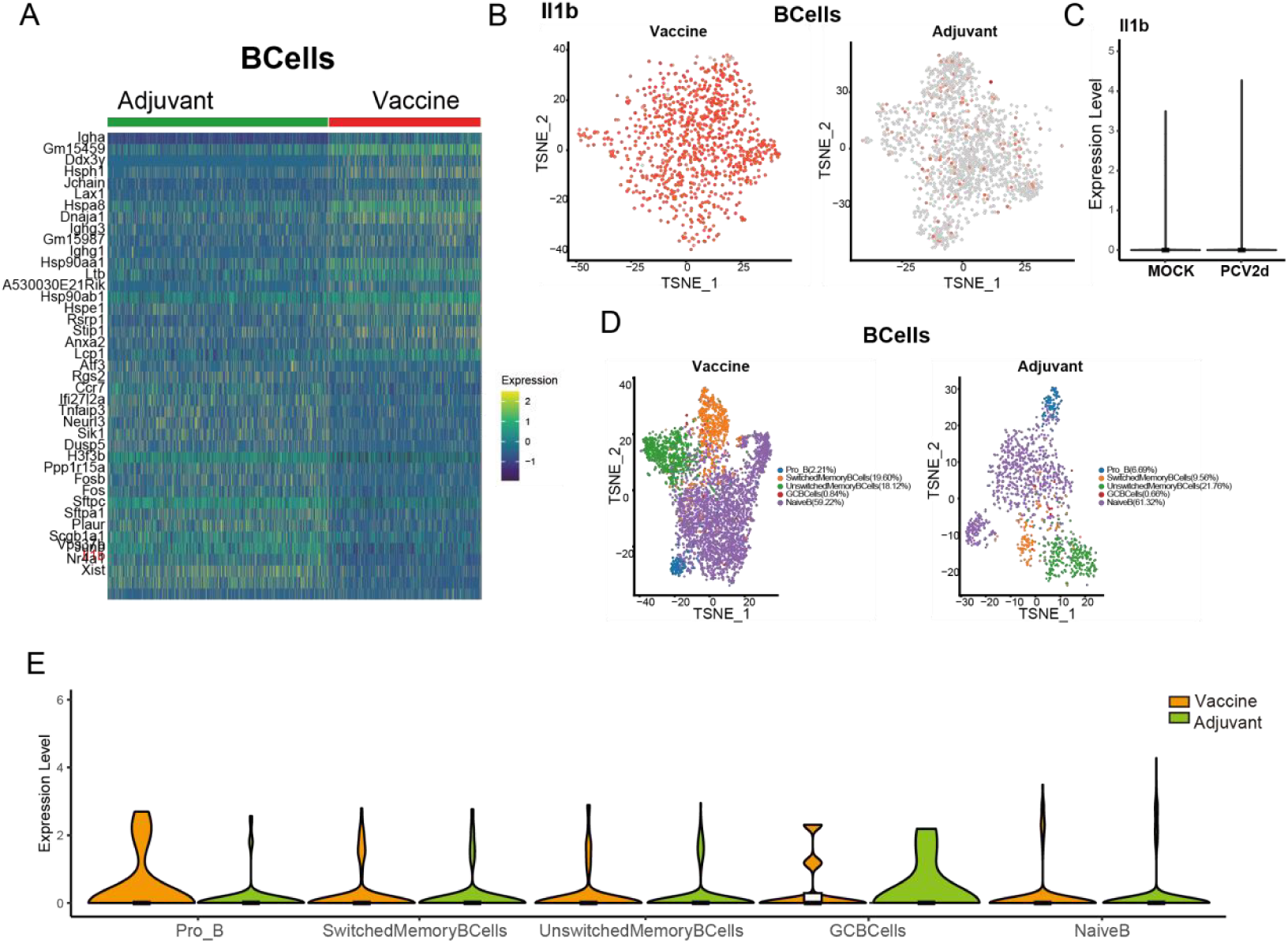
Analysis of B cells in Vaccine group and Adjuvant group. (A) Total differential genes in Bcells under two treatments. (B) IL-1b expression in B-cells of both treatment groups. (C) IL-1b gene violin expression of cello in Bcells under two treatment groups. (D) Secondary grouping of Bcells under two treatment groups. (E) IL-1b violin expression in two subsets of B cells.

**Supplementary Figure 6:**
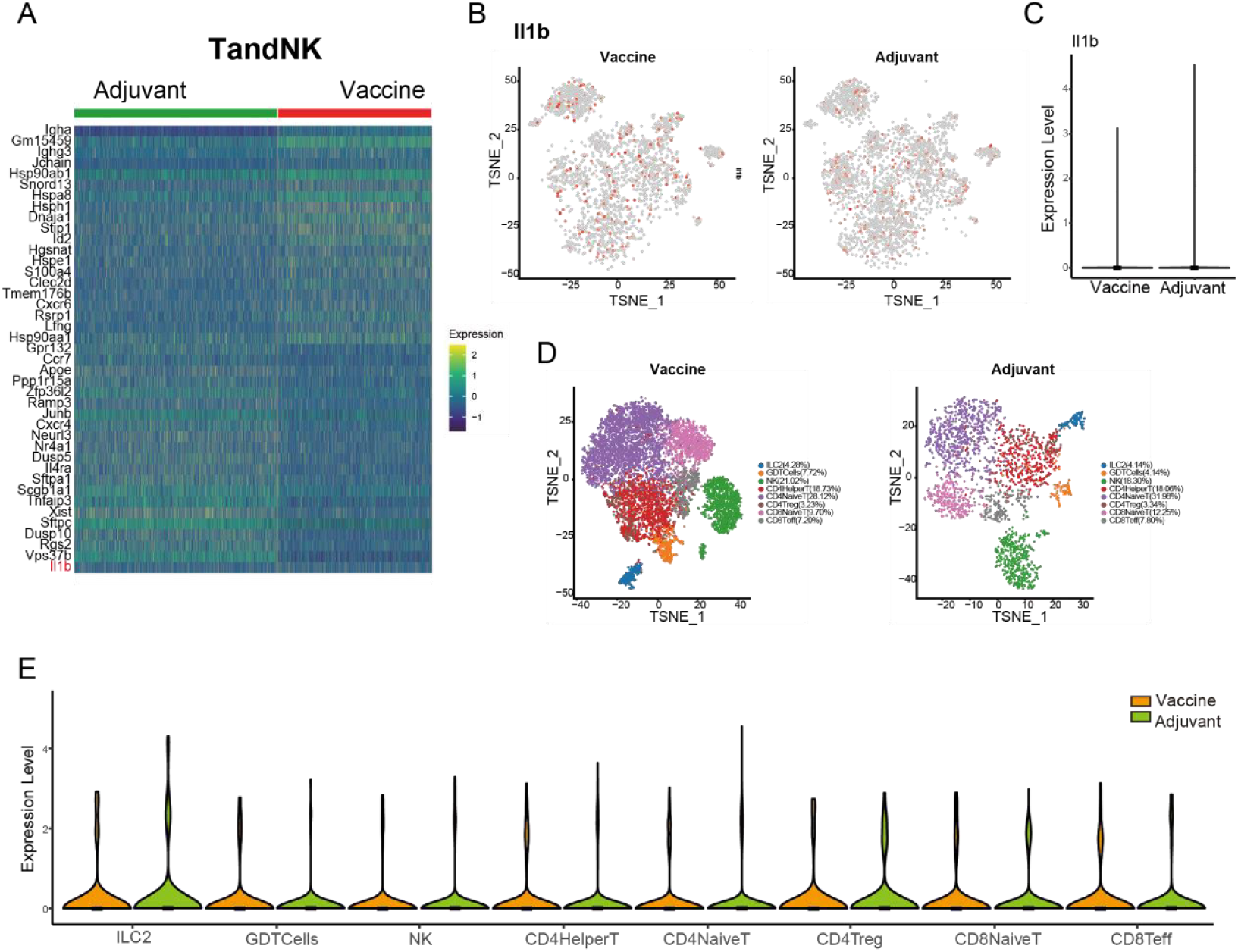
TandNK analysis chart of Vaccine group and Adjuvant group. (A) TandNK total differential genes under two treatments. (B) IL1b expression in TandNK class of two treatment groups. (C) IL1b gene violin expression level of TandNK class under the two treatment groups. (D) TandNK was divided into two groups. (E) Violin expression of IL-1b in T and NK subpopulations.

**Supplementary Figure 7:**
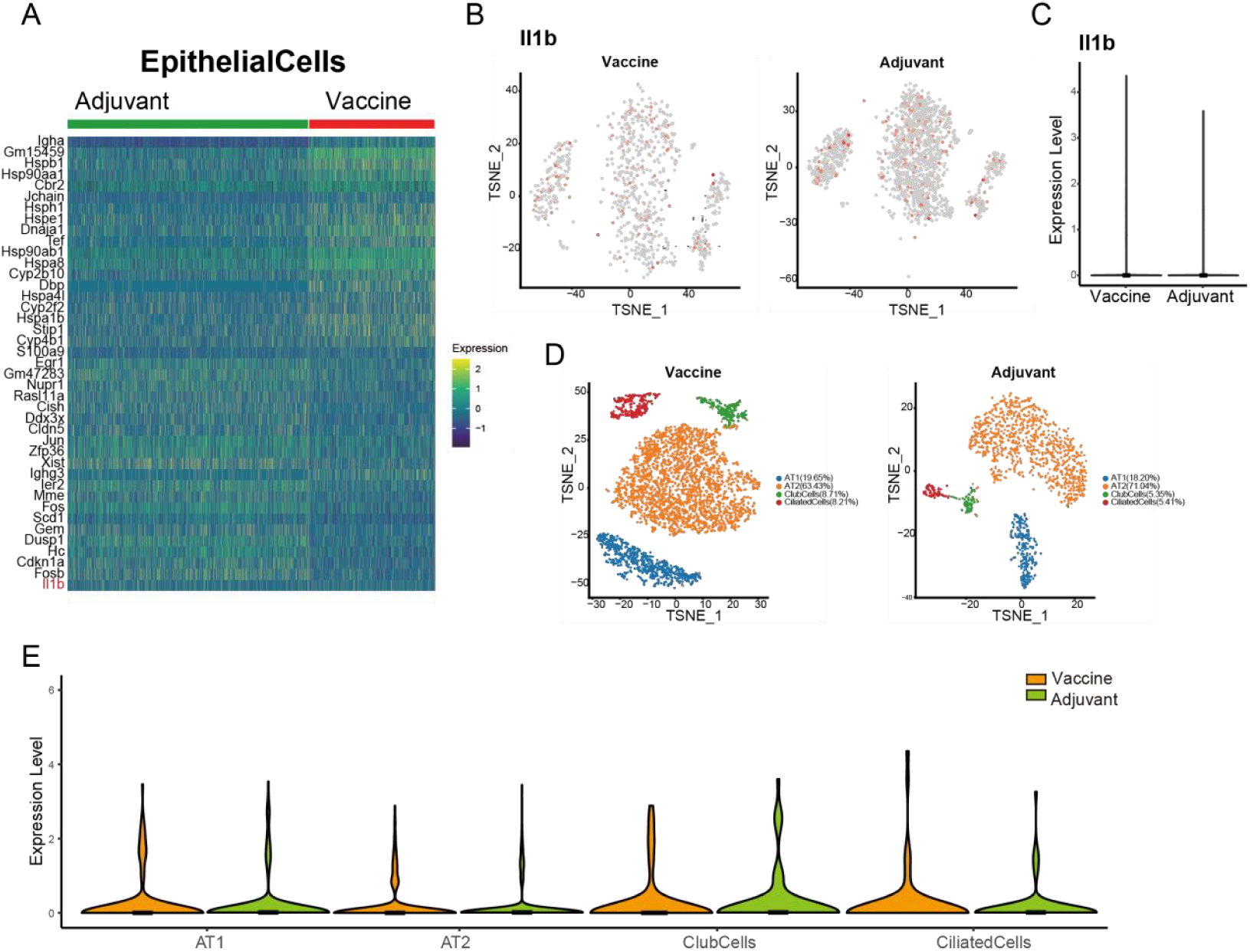
EpithelialCells analysis of Vaccine group and Adjuvant group. (A) Total differential genes in EpithelialCells under two treatments. (B) IL1b expression in EpithelialCells of both treatment groups. (C) IL1b gene violin expression in EpithelialCells under both treatment groups. (D) Subgrouping EpithelialCells under two treatment groups. (E) Violin expression of IL-1b in two subgroups of EpithelialCells.

**Supplementary Figure 8:**
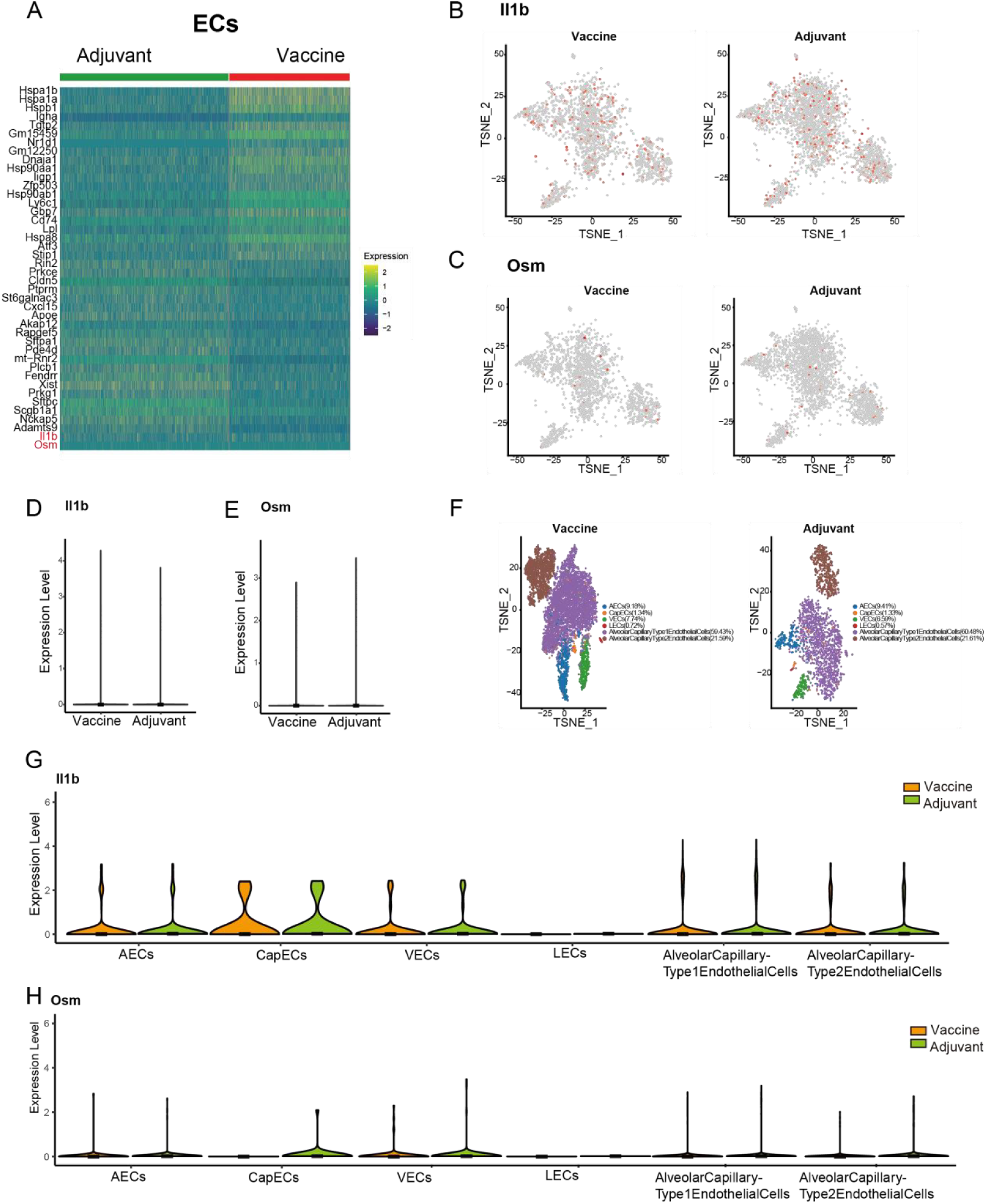
ECs analysis chart of Vaccine group and Adjuvant group. (A) Total differential genes in ECs under two treatments. (B) IL1b expression in ECs of both groups. (C) Osm expression in ECs of two treatment groups. (D) Violin expression of IL1b gene in ECs under both treatment groups. (E) Violin expression of Osm gene in ECs under two treatment groups. (F) Secondary clustering of ECs under two treatment groups. (G) Violin expression of IL-1b in ECs (H) Violin expression of Osm in ECs.

